# A side-by-side comparison of peptide-delivered antisense antibiotics employing different nucleotide mimics

**DOI:** 10.1101/2023.07.11.548539

**Authors:** Chandradhish Ghosh, Linda Popella, V. Dhamodharan, Jakob Jung, Lars Barquist, Claudia Höbartner, Jörg Vogel

## Abstract

Antisense oligomer (ASO)-based antibiotics that target mRNAs of essential bacterial genes have great potential for counteracting antimicrobial resistance and for precision microbiome editing. To date, the development of such antisense antibiotics has primarily focused on using phosphorodiamidate morpholino (PMO) and peptide nucleic acid (PNA) backbones, largely ignoring the growing number of chemical modalities that have spurred the success of ASO-based human therapy. Here, we directly compare the activities of seven chemically distinct 10mer ASOs, all designed to target the essential gene *acpP* upon delivery with a KFF-peptide carrier into *Salmonella.* Our systematic analysis of PNA, PMO, phosphorothioate-modified DNA (PTO), 2’-methylated RNA (RNA-OMe), 2’-methoxyethylated RNA (RNA-MOE), 2’-fluorinated RNA (RNA-F) and 2’-4’-locked RNA (LNA) is based on a variety of *in vitro* and *in vivo* methods to evaluate ASO uptake, target pairing and inhibition of bacterial growth. Our data show that only PNA and PMO are efficiently delivered by the KFF peptide into *Salmonella* to inhibit bacterial growth. Nevertheless, the strong target binding affinity and *in vitro* translational repression activity of LNA and RNA-MOE make them promising modalities for antisense antibiotics that will require the identification of an effective carrier.

## INTRODUCTION

Microbial resistance to frontline drugs has necessitated the search for alternatives to conventional antibiotics. Short single-stranded antisense oligomers (ASOs) that inhibit the translation of essential bacterial genes are promising candidates (1,2). They have been shown to be effective antibacterials against a range of pathogens (3–8). Their efficacy has also been validated in animal models of infection with *Escherichia coli*, *Pseudomonas aeruginosa* and *Klebsiella pneumoniae* either as monotherapy or in combinations with other antibiotics (9–12).

Antisense antibiotics have potential advantages over conventional antibiotics. First, given their sequence-specific mode of action, they can be designed to be species-specific, thus allowing selective targeting of pathogenic bacteria over commensals (1,13,14). Second, development of resistance due to mutations in the target mRNA can be reversed by changing the ASO sequence (15). Third, ASOs can in principle target any transcript of interest and can therefore also be used to reinstate the sensitivity of drug-resistant bacteria to antibiotics (10,16). Nevertheless, despite their success in laboratory settings, no antisense antibiotic has been approved for clinical use. For antisense antibiotics to progress to clinical development, improvements in their design and delivery are needed.

Antibacterial ASOs are generally modular, comprising an antisense effector and a delivery vehicle. The antisense moiety is typically 10-15 nucleotides in length and designed to hybridize to the ribosome binding site (RBS) or start codon of an mRNA of interest. It sterically blocks ribosome binding and thereby prevents translation of the target protein (17). Since ASOs cannot be directly taken up by bacteria, they require vehicles such as cell-penetrating peptides (CPPs) to deliver them across the bacterial envelope into the cytosol, their place of action. While there has been much effort to diversify the delivery strategies within and beyond the realms of CPPs (18–20), less attention has been paid to finding the best chemistry for the antisense moiety.

Due to the rapid degradation of unmodified RNA by nucleases, the antisense moiety requires a modified nucleic acid or mimetic with improved stability and nuclease resistance. To date, peptide nucleic acid (PNA) and phosphorodiamidate morpholino (PMO) have been the most popular modalities for the use as antisense antibiotics (21). In PNAs, the phosphodiester backbone of the nucleic acid is replaced by a pseudo-peptide backbone, while in PMOs the five-membered ribose sugar is replaced by a six-membered morpholine ring. Both modalities are neutral in charge, which improves their target affinity and eases delivery by charged CPPs.

Efforts in medicinal chemistry have led to a large range of additional modifications that enable ASO optimization in terms of pharmacokinetics, pharmacodynamics and bioavailability. These have been applied to oligonucleotide drug design in eukaryotic systems and several oligonucleotides have been approved for clinical use against a range of human diseases such as retinitis, macular degeneration, hypercholesterolemia, amyloidosis and muscular dystrophy (22). Successful ASO modalities include modifications on the 2′-hydroxy group on the ribose sugar of a nucleotide, such as 2′-4′ carbon bridged nucleic acids (BNA and LNA), 2’-methylated RNA (RNA-OMe), 2ʹ-*O*-methoxyethyl (2ʹ-MOE) and 2ʹ-Fluoro (2ʹ-F). Another common modification is the substitution of the non-bridging oxygen in the phosphate backbone of nucleic acids, for example phosphorothioate-modified DNA (PTO) (23–28) (Figure 1A). As it stands, few studies have explored the antibacterial potential of these ASO modalities. 2ʹ-MOE and 2ʹ-F have not been tested in bacteria at all. Moreover, there has been no side-by-side comparison of the efficacy of these different ASO chemistries in terms of target binding, inhibition of target gene translation and their antibacterial activity.

**Figure 1.**
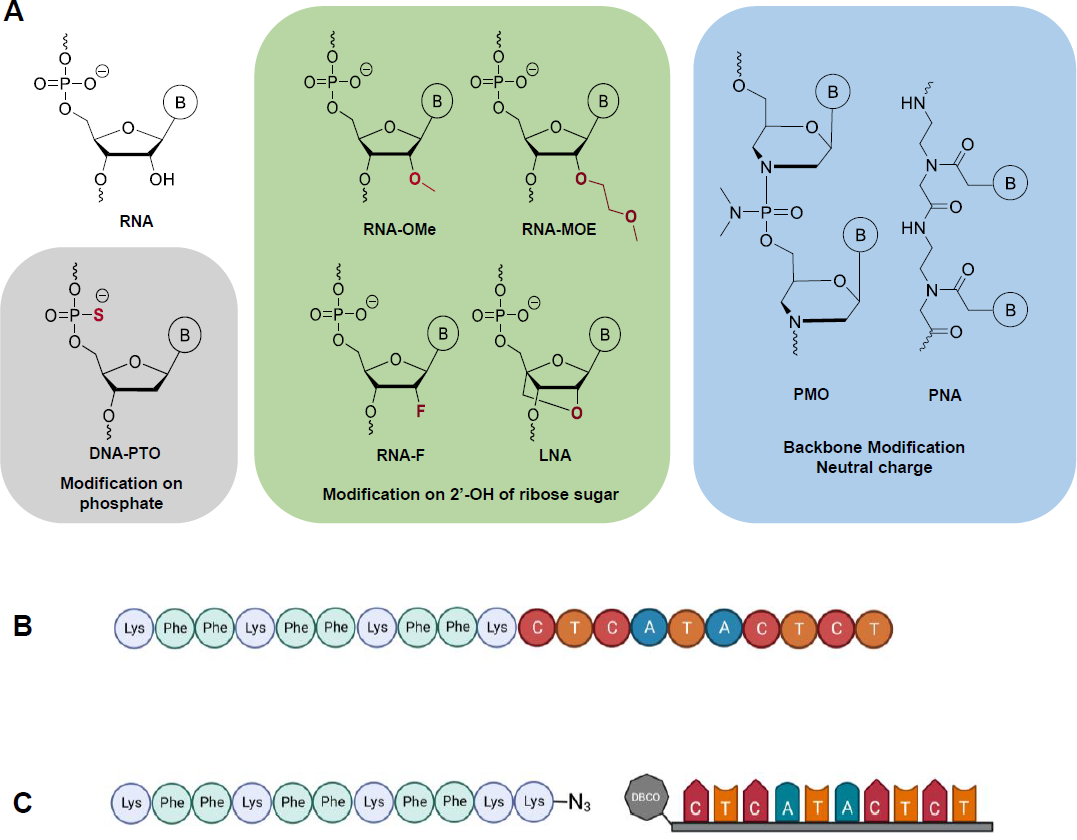
Design and synthesis of the different ASOs used in the study. (A) The different chemical modalities used in this study can be grouped into three different categories: Modifications of the phosphate backbone (grey), modifications of the ribose sugar (green) and alternative backbone chemistries (blue). The grey and the green boxes comprise ASOs that bear net negative charges; ASOs in the blue box are charge-neutral. (B) The KFF-PNA conjugates were synthesized as fusion peptides on a peptide synthesizer. (C) Schematic for the strain promoted alkyne-azide click reaction between the peptide (synthesized on a peptide synthesizer) and the ASO (synthesized on an oligonucleotide synthesizer). Created with BioRender.com.

Over the years, methodology has been developed to study the uptake, target binding and specificity of ASO antibiotics. For example, the Nielsen lab has established methods to study the internalization of peptide-PNA conjugates in bacteria (29). Our own efforts have been directed at using RNA sequencing (RNA-seq) to read out how ASOs affect target mRNA levels in bacteria and how bacteria respond to ASO treatment in general (30,31). This is complemented by the development of assays and algorithms to study ASO potency and specificity *in vitro* and *in vivo* (31–33). Here, using these tools, we have systematically compared seven ASO modalities in their antibacterial efficacy. We compared their binding affinity and inhibition of target protein synthesis in a cell-free system and, upon conjugation to the commonly used carrier peptide (KFF)_3_K (henceforth, KFF), we assessed their effect on the growth and physiology of the model pathogen *Salmonella enterica* (henceforth, *Salmonella*). Additionally, we studied the delivery of these ASOs across *Salmonella* membranes. Our results suggest that although LNA and RNA-MOE tightly bind and efficiently inhibit target mRNA translation *in vitro*, the KFF carrier peptide is unable to deliver these negatively charged ASOs (negASOs) into the bacteria. This reinforces the use of neutral backbone ASOs for KFF-mediated delivery, yet indicates that negASOs might be efficient antisense antibacterials when delivered with a compatible carrier.

## MATERIAL AND METHODS

### Oligomers, reagents and bacterial strains

The antisense oligonucleotides were purchased from Biomers.net GmbH (Ulm, Germany). PNAs, peptides and peptide-conjugated PNAs (PPNAs) were obtained from Peps4LS GmbH (Heidelberg, Germany). Quality and purity of these constructs were verified by mass spectrometry and HPLC. ASOs, peptides and PPNAs (Table 1) were dissolved in water and heated at 55 °C for 5 minutes (min) before determining the concentration by using a NanoDrop spectrophotometer (A _260 nm_ for ASO and KFF-ASO, A_205 nm_ for peptides). Aliquots of ASOs and peptides were stored at - 20 °C and heated at 55 °C for 5 min before preparing working dilutions. Low retention pipette tips and low binding Eppendorf tubes (Sarstedt) were used throughout. *Salmonella enterica* serovar Typhimurium strain SL1344 (provided by D. Bumann, Biocenter Basel, Switzerland; internal strain number JVS-1574) was used in this study. Bacteria were cultured in non-cation adjusted Mueller-Hinton Broth (MHB, BD Difco^TM^, Thermo Fisher Scientific) with aeration at 37°C and 220 rpm shaking.

**Table 1.** The different chemical modalities used in this study and their physical parameters.

| Chemical entity | Net charge | Molecular weight | Binding affinity (EC<sub>50</sub>, nM)* |
| --- | --- | --- | --- |
| RNA-F | -10 | 3540.55 | 72.8 ± 36.2 |
| RNA-OMe | -10 | 3660.75 | 4.0 ± 0.2 |
| RNA-MOE | -10 | 4101.02 | 0.3 ± 0.2 |
| LNA | -10 | 3752.72 | 0.3 ± 0.1 |
| DNA-PTO | -10 | 3576.48 | Could not be determined |
| PMO | 0 | 3936.5 | 5.9 ± 2.7 |
| PNA | 0 | 2579.02 | 0.4 ± 0.4 |
| (KFF)<sub>3</sub>KK(N<sub>3</sub>) | +5 | 1565.90 | N.A. |
| (KFF)<sub>3</sub>K | +5 | 1427.79 | N.A. |

\*Mean ± Standard deviation (two independent experiments). N.A. stands for Not applicable.

### Electrophoretic mobility shift assay

The electrophoretic mobility shift (EMSA) assay was performed with a Cy5 5’-labelled RNA (purchased from Eurofins) encompassing the sequence spanning nucleotides −20 to +20 of *acpP* relative to the translation start codon. For annealing, TM buffer (10 mM Tris-HCl, 50 mM MgCl_2_) was used. First, 20 µM RNA was denatured by incubation at 95 °C for 5 min followed by placement on ice. Then 1 µL of RNA was added to PCR tubes containing 8 µL of water. To this 1 µL of ASOs at the indicated concentrations (25 µM or 50 µM) were added to adjust ASO:RNA ratios of 1.25:1 and 2.5:1. As untreated negative control water was added instead of ASO, while a scrambled PNA sequence served as sequence-unspecific negative control (Figure 2 and Figure S2). These mixtures were incubated for 30 min at room temperature (RT), followed by the addition of 10 µL 2x Gel Loading (GL) buffer for subsequent loading on native 15 % PAA RNA gel without urea (20 µL/sample; 6 µL of a 50 bp DNA marker + 14 µL 1x GL). The gel was run at 100-120 V for 2-3 h, concomitantly stained with ethidium bromide for 15 min and imaged on Geldoc (Isogen Life Science B.V., De Meern, the Netherlands). Figure 2 shows a representative image out of three replicates.

**Figure 2.**
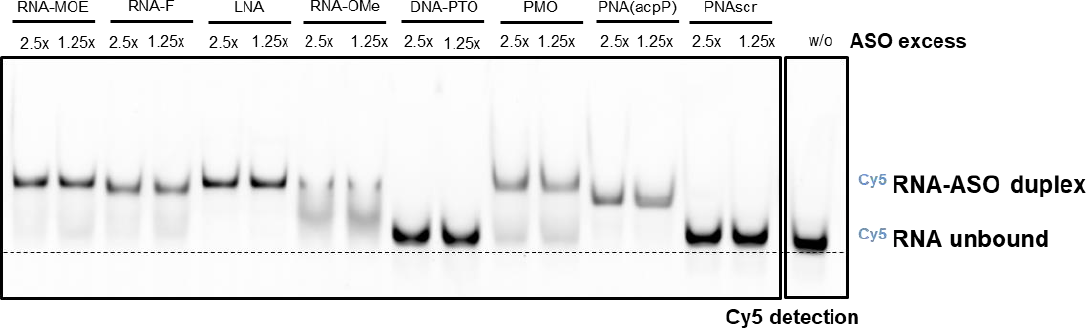
Electrophoretic mobility shift assay. Retardation of electrophoretic mobility of Cy5 5’-labelled *acpP* target RNA (40 nt) due to ASO hybridization as observed in native PAGE. The black dashed line indicates the migration pattern of unbound RNA. A representative example of three independent experiments is shown.

### Determination of binding affinities using Microscale thermophoresis (MST)

For each binding experiment, Cy5 5’-labelled RNA (as described above) was diluted to 10 nM in Buffer A (50 mM Tris-HCl pH 7.6, 250 mM KCl, 5 mM MgCl2, 1 mM DTT, 5% glycerol supplemented with 0.05% Tween 20). A series of 16 tubes with different ASO dilutions (500 nM to 0.015 nM) were prepared in Buffer A. For measurements, each ASO dilution was mixed with one volume of labeled RNA, which led to a final concentration of 5.0 nM labeled RNA and 250 nM to 0.007 nM of ASOs. The reaction was mixed by pipetting and incubated for 30 min at RT in the dark. Capillary forces were used to load the samples into Monolith NT.115 Premium Capillaries; measurements were performed using a Monolith Pico instrument (NanoTemper Technologies Gmbh, Munich, Germany) at an ambient temperature of 25 °C. Instrument parameters were adjusted to 5% LED power, medium MST power, and MST on-time of 2.5 s. An initial fluorescence scan was performed across the capillaries to determine sample quality, which was followed by 16 thermophoresis measurements. Measurements that showed fluorescence inhomogeneity or aggregation were excluded. Data of two independent measurements were analyzed for the ΔFnorm value determined by the MO Affinity Analysis software (NanoTemper Technologies Gmbh, Munich, Germany). The fitting for EC_50_ values was done using the Hill equation within the MO Affinity Analysis software. Data are presented as the mean EC_50_ values of two independent experiments.

### Synthesis of target RNA using in vitro transcription

For the generation of target RNA::*gfp* fusion constructs, we used a previously established protocol (31) and the oligonucleotides JVO-19760/JVO-19761 (Supplementary Table 1) to amplify the genomic region spanning nucleotides –40 to + 51 relative to the translational start codon of *acpP*. The sense oligonucleotide was fused to a T7 promoter sequence for subsequent transcription of the final fusion construct *in vitro*. The antisense oligonucleotide contained a 30 nt overlapping region with *gfp* for subsequent fusion PCR with full length *gfp* (pXG-10; JVO-19762/JVO-19763) (Supplementary Table 1). The PCR products were purified with NucleoSpin Gel and PCR Clean-up (Macherey-Nagel) according to the manufacturer’s instructions and DNA concentration was quantified with a NanoDrop spectrophotometer (*A*_260 nm_).

PCR products were used for T7 RNA polymerase-driven transcription *in vitro* using the MEGAscript T7 kit (Ambion/Thermo Scientific), according to the manufacturer’s instructions and analogous to our previous publication (31). The concentration of the RNA solution was quantified with Qubit (Fisher Scientific) and the expected product size and RNA integrity was verified on a 6% PAA, 7 M urea gel.

### In vitro translation and western blotting

For *in vitro* translation of target RNA::*gfp* fusion constructs, the PURExpress® In Vitro Protein Synthesis Kit (New England Biolabs, E6800L) was used as described previously (31). Briefly, the final volume of each *in vitro* translation reaction was set to 10 µL. 1µL of different concentrations of ASOs ranging from 10 µM to 0.625 µM were added to 1 µL of *in vitro* transcribed heat-denatured RNA (1µM) to adjust the final RNA concentration to 100 nM. The mixtures were pre-incubated at 37 °C for 5 min. Then 4 µL of Solution A and 3 µL of Solution B were added to the pre-annealed mixtures. After 2 h at 37 °C, protein samples were denatured in 1x reducing protein loading buffer (62.6 mM Tris–HCl pH 6.8, 2 % SDS, 0.1 mg/mL bromphenol blue, 15.4 mg/mL DTT, 10 % glycerol) at 95 °C for 5 min.

To visualise *in vitro* translated proteins, samples were separated on SDS-PAA (12 % PAA) gels, with subsequent semi-dry western blot transfer on polyvinylidene fluoride (PVDF) membranes. Membranes were blocked with 10% skim milk (in 1x TBS-T) for 1 h and probed with anti-GFP antibody (1:1,000; Sigma-Aldrich) in 5 % skim milk (in 1x TBS-T) overnight at 4 °C. After washing in 1x TBS-T and incubation with an HRP-conjugated secondary antibody (1:5,000; ThermoScientific) in 1x TBS-T, the membrane was incubated with developing solution kit (Amersham™ ECL Select™ Western Blotting Detection Reagent) and protein levels were detected using an ImageQuant LAS 500 (GE Healthcare Life Sciences). Images were processed and band intensities quantified using ImageJ. Protein bands were normalized to the water control.

### Synthesis of ASO-peptide conjugates

Due to their pseudo-peptide backbone, KFF-PNA conjugates can readily be synthesised as a fusion peptide using a peptide synthesizer. All other ASOs are synthesized on an oligonucleotide synthesizer with an appended 5’ terminal dibenzocyclooctyne (DBCO) moiety, which enabled strain-promoted alkyne-azide cycloadditions (SPAAC) (copper-free click chemistry) with azides (34). To facilitate coupling to KFF, a terminal lysine whose ε-amine was replaced by an azide was added as the C-terminal amino acid (sequence: (KFF)_3_KK(N_3_)).

All ASOs with a DBCO terminated linker (Figure 1B) were purchased from Biomers (Ulm, Germany) with the exception of PMO, which was purchased from GeneTools, LLC (Philomath, Oregon, USA). The DBCO terminated linker was not available from GeneTools, LLC (Philomath, Oregon, USA), so we used PMO terminating with a cyclooctyne linker. Of note, the antisense PMO-peptide conjugates previously reported to have antibacterial activity did not use click chemistry for conjugation to the peptides but use amide coupling chemistry (35,36).

In a typical reaction, 1 equivalent of (in 50 µL water) of ASO with the DBCO moiety was added to 3 equivalents of (KFF)_3_KK(N_3_) (in 450 µL water). The reaction was carried out at 4 °C overnight. Precipitation was observed for some of the compounds (e.g. LNA, DNA-PTO, RNA-MOE) when mixed with the (KFF)_3_KK(N_3_) peptide, which was usually resolved by adding equal volumes of acetonitrile or formamide. The ASO-CPP conjugate was separated from the reactants by running them on 20 % polyacrylamide gel, excising the appropriate bands and extracting them into TEN buffer. The compounds were then purified using size exclusion chromatography (Äkta pure, Cytiva). Each ASO-CPP conjugate was characterized using MALDI mass spectrometry (Supplementary Figure S5). The samples were stored as lyophilized powders at −20 ⁰C for further usage.

### Salmonella growth assays

An overnight bacterial cell culture was diluted 1:100 in fresh MHB and grown to OD_600_ 0.5. The obtained culture was diluted to approximately 10^5^ cfu/ml in non-cation-adjusted MHB. Subsequently, 190 µl bacterial solution was dispensed into a 96-well plate (Thermo Fisher Scientific) along with 10 µl of a 200 µM solution of the ASOs. Growth was monitored by measuring the OD at 600 nm every 20 min in a Synergy H1 plate reader (Biotek) with continuous double-orbital shaking (237 cpm) at 37 °C for 24 h.

### RNA isolation for sequencing

RNA isolation was performed using previously published protocols (30,31). Bacterial overnight cultures were diluted 1:100 in fresh MHB and grown to OD_600_ 0.5. The cultures were diluted to approximately 10^6^ cfu/ml in non-cation-adjusted MHB. Afterwards, an aliquot of the bacterial solution was transferred into 5 ml low-binding tubes (LABsolute) containing the ASOs solution (adjusted such that the final concentration was 5 μM for all tested compounds). As a negative control, cells were treated with the respective volume of sterile nuclease-free water, which was used as solvent for the ASOs. After incubating the samples for 15 min at 37 °C, RNAprotect Bacteria (Qiagen) was added according to the manufacturer’s instructions. Following a 10-min incubation, cells were pelleted at 4 °C and 21,100 × g for 20 min. The supernatant was discarded and pellets were either directly used or stored at –20°C (<1 day) for subsequent bacterial RNA isolation.

Total RNA was purified from bacterial pellets using the miRNeasy Mini kit (Qiagen) according to protocol #3 described in (30). Briefly, cells were resuspended in 0.5 mg/ml lysozyme (Roth) in TE buffer (pH 8.0) and incubated for 5 min. Afterwards, RLT buffer supplemented with β-mercaptoethanol, and ethanol were added according to the manufacturer’s instructions. After sample loading, column wash-steps were performed according to the manual. RNA concentration was measured with a NanoDrop spectrophotometer.

### RNA-seq

For transcriptomic analyses, RNA samples were processed and subjected to RNA-seq at Core Unit SysMed (University of Würzburg, Germany). Sequencing was run using a previously published protocol (31). Briefly, RNA quality was checked using a 2100 Bioanalyzer with the RNA 6000 Pico/ Nano kit (Agilent Technologies). RNA samples were DNase-treated using DNAse I kit (Thermo Fisher), followed by ribosomal RNA depletion using Lexogen’s RiboCop META rRNA Depletion Kit protocol according to manufacturer’s recommendation. Subsequently, cDNA libraries suitable for sequencing were prepared using CORALL Total RNA-Seq Library Prep protocol (Lexogen) according to manufacturer’s recommendation with 14–25 PCR cycles. Library quality was checked using a 2100 Bioanalyzer with the DNA High Sensitivity kit (Agilent Technologies). Sequencing of pooled libraries, spiked with 5% PhiX control library, was performed at 10 million reads/sample in single-end mode with 75 nt read length on the NextSeq 500 platform (Illumina) using High output sequencing kits. Demultiplexed FASTQ files were generated with bcl2fastq2 v2.20.0.422 (Illumina).

### RNA-seq data analysis

RNA-seq data analysis was performed similarly to our previous analysis of PNA activity in *S. enterica* (30). Briefly, adapters and low-quality bases (Phred quality score <10) of raw reads were trimmed using BBduk. Then the RNA-seq reads were mapped to the the *S. enterica* serovar Typhimurium reference genome and the three plasmids pSLT SL1344, pCol1B9 SL1344, and pRSF1010 SL1344 (37) using BBMap (v38.18) and genomic features were assigned using the featureCounts command of the Subread (2.0.1) package (38). For the coverage plots in Figure 8, the bamCoverage (v3.3.2) method of the deepTools platform was applied (39).

### Normalization and differential expression analysis

Downstream RNA-seq analysis was performed using packages from the R/Bioconductor project. Raw read counts of all conditions were imported into edgeR (v3.34.1) (40) and analysed for differential expression. To filter low-count reads, features with less than 1.38 counts per million (CPM) in at least 5 libraries were ignored for further analysis. The cutoff of 1.38 CPM was set to 10/L, with L being the minimum library size across all samples in millions, as described in (41). Libraries were normalized using edgeR’s trimmed mean of *M* values (TMM) normalization (40). The glmFit function was applied to estimate quasi-likelihood dispersions and contrasts for differential expression analysis were tested using the glmQLTest function. Features with an absolute fold change >2 and false discovery rate (FDR) (42) adjusted *P*-values <0.001 were considered significantly differentially expressed. The results were plotted as heatmaps using the ComplexHeatmap (2.12.0) package (43).

### KEGG pathway analysis

To perform gene-set analysis, features were assigned to KEGG pathways (44) with the R package KEGGREST (v1.32.0). Additionally, genes were assigned to gene sets of regulons curated in (45). To assess enriched KEGG pathways and regulons, rotation gene set testing was performed using the FRY function of the ROAST package(46). Top up- or downregulated gene sets covering >10 genes and an FDR-adjusted p-value <0.01 (marked with an asterisk) in at least one condition were visualized in Figure 7. The colour in the heatmap denotes the median log_2_FC of the respective gene set. The figure shows gene sets ranking among the top 10 significant gene sets (lowest P-value) in at least one sample.

### Quantification of intracellular AcpP protein levels post ASO treatment

*Salmonella* wild type (WT) and SL1344 AcpP::3xFLAG (chromosmal fusion of full length *acpP* to 3XFLAG-tag as described earlier(47); internal strain number JVS-12398) were grown overnight in non-cation adjusted MHB. Bacterial cultures were diluted 1:100, grown to OD_600_ 0.5 and subsequently diluted to approximately 10^6^ cfu/ml in MHB. Then, 1.9 ml of each diluted bacterial solution was transferred into 5 ml low-binding tubes, pre-incubated at 37 °C for 10 min and treated with 100 µl of ASOs to adjust a final concentration of 5 µM. Water was added as negative control. The treatment with PNA and PMO was performed with aeration at 37 °C and 220 rpm shaking for 30, 60, 120 and 240 minutes while for all other ASOs, they were treated for 120 min only. Subsequently, the cells were collected by centrifugation at 4 °C and 16,000 ×g for 5 min, resuspended in 1× Protein Loading Buffer (62.6 mM Tris–HCl pH 6.8, 2 % SDS, 0.1 mg/ml bromophenol blue, 15.4 mg/ml DTT, 10 % glycerol) and denatured at 95 °C for 10 min.

For detection of AcpP::3xFLAG protein levels, samples were separated on an 12% SDS-PAA gel and transferred onto a Nitrocellulose membrane via semi dry western blotting. The membrane was probed with an α-FLAG antibody (Sigma; 1:1,000 in 1x TBS-T containing 3% BSA) at 4 °C overnight. After washing in 1x TBS-T, the membrane was incubated with a secondary HRP-conjugated anti-mouse antibody (Thermo Scientific; 1:5,000 in 1x TBS-T containing 1% BSA) at RT for 1 hour. Excessive antibody was removed by repeated wash steps in 1x TBS-T and the membrane was developed by Amersham™ ECL Select™ Western Blotting Detection Reagent Kit (Cytiva).

### Outer membrane permeabilization

The experiments were performed by modifying previously published protocols (48–50). *Salmonella* (WT SL1344, JVS-1574) were grown overnight in 4 ml MHB medium at 37 °C (with shaking at 220 rpm). Then the culture was diluted 1:100 and grown until OD 0.5 (∼2×10^8^ cfu/mL). This culture was harvested and washed two times with 5mM HEPES and further diluted to OD 0.2 in 5 mM HEPES supplemented with 5 mM Glucose (pH 7.2). These cells were then incubated with *N*-phenylnaphthylamine (NPN) (Sigma, USA) at a final concentration of 10 µM. 190 µl of NPN treated cells were transferred into 96-well back plates with transparent bottom and fluorescence (excitation = 350 nm, emission = 420 nm) was recorded for 60 min (reading was performed every minute) using a Synergy H1 plate reader (Biotek). On the fourteenth minute, 10 µl of the test compounds (10 µM final concentration) was added into the wells. CTAB (Sigma, USA) 200 µM or Polymyxin B (Sigma, USA) 10 µg/ml (final concentration of 0.5 µg/ml) were used as positive controls, while water was used as negative control. The fluorescence of the wells were monitored using the same settings as above. The experiment was performed three times and Figure 10 is based on a single representative experiment.

### MALDI-TOF based analysis of ASO-peptide conjugate uptake into bacterial cells

The experiments were performed by modifying previously published protocols (29). From an overnight culture of *Salmonella*, a second culture was inoculated (1:100) and grown to OD_600_ of 0.5 (corresponding to ∼2×10^8^ cfu/mL). These bacterial cells was washed twice in MHB and resuspended in 95 µl of MHB with an additional incubation at 37°C for 5-10 min. Sterile ASO-peptide conjugates were prepared at concentrations of 200 µM. 5 µl of the 200 µM working solutions were added to 95 µl of bacteria in culture media, leading to a final concentration of the compounds of 10 µM. 5 µl of H_2_O was added as a negative control. The tubes were incubated at 37°C for 15 min with shaking at 300 rpm. Subsequently, they were centrifuged at 8,000 *x* g for 5 min at 4 °C. The supernatant was aspirated and the pellet was washed with 0.01 M Tris-HCl before being resuspended in 200 µl of 0.01M Tris-HCl containing 0.5 M Sucrose, 150 µg/ml lysozyme, 10 mM EDTA and incubated at 37 °C overnight. These tubes were then centrifuged at 2,000 *x* g at 4°C for 10 min. The supernatant obtained represents the periplasm. The pellet containing spheroplasts was gently washed with 0.01 M Tris-HCl containing 0.5 M Sucrose and resuspended in 20 ml of B-PER™ Bacterial Protein Extraction Reagent (Thermo Fischer Scientific) for 1 h. After this, the spheroplasts were subjected to sonication (20kHz for 2×10 s), incubated on ice for 5 min and centrifuged at 20,000 *x* g for 30 min at 4 °C. The supernatant represents the cytoplasm. Both the periplasm and cytoplasm fragments were then mixed with 2, 6-Dihydroxyacetophenone (DHAP) in ACN 1:1, spotted on a MALDI plate and analyzed for MALDI-TOF-MS in a Bruker DaltonicsUltrafleXtreme Instrument. The experiment was performed two times and data of one representative experiment are shown.

## RESULTS

### Study design

For our side-by-side comparison of different ASO modalities, we selected three representatives of backbone-modified synthetic nucleotides (PNA, PMO, and PTO) and four ribose-modified nucleotides (2ʹ-OMe, 2ʹ-MOE, 2ʹ-F, and LNA); members of both groups are successfully used as ASO therapeutics for human diseases (22). PNA and PMO are charge-neutral, while all the other ASOs mentioned above are negatively charged due to their phosphate backbone (Figures 1A and S1, Table 2). We choose to target the essential gene *acpP* (acyl carrier protein), since PNA-mediated silencing of *acpP* protein synthesis rapidly and effectively inhibits *Salmonella* growth (30). In addition, the concomitant and selective depletion of *acpP* mRNA by an as yet unknown mechanism can be read out by RT-qPCR, northern blot or RNA-seq (51). Each of the seven different ASOs was designed to be complementary to nucleotides −5 to +5 relative to the translational start site of *acpP* (sequence of the ASOs: CTCATACTCT). We considered a 10mer ASO optimal to balance hybridization strength and efficient cellular uptake, since increasing ASO length hinders internalization (52,53).

**Table 2.** MS-based detection of ASO-peptide fragments in *Salmonella* periplasm and cytoplasm 15 min post treatment.

| **KFF-ASO** | **Periplasm** | **Fragments detected** |
| --- | --- | --- |
| (KFF)<sub>3</sub>K-PNA | ✓ | PNA (+KFF) |
| (KFF)<sub>3</sub>KK(N<sub>3</sub>)-PMO | ✓ | PMO |
| (KFF)<sub>3</sub>KK(N<sub>3</sub>)-LNA | N.D. | - |
| (KFF)<sub>3</sub>KK(N<sub>3</sub>)-RNA-F | N.D. | - |
| (KFF)<sub>3</sub>KK(N<sub>3</sub>) RNAMOE | N.D. | - |
| (KFF)<sub>3</sub>KK(N<sub>3</sub>)RNA-OMe | N.D. | - |

N.D. stands for Not detected

For experiments involving delivery into *Salmonella*, we fused the different ASOs to the KFF carrier peptide (Figures 1B-C). Although the covalent conjugation of the cationic amphiphilic KFF peptide and the anionic ASOs might lead to the formation of aggregates due to charge-charge interactions, we chose KFF for three reasons. First, KFF has previously been used as carrier for different ASO modalities (such as PNA, PMO and LNA) and has been successfully applied to different bacterial species (3,6,23,54). Second, in a mini-library of different CPP-ASOs, KFF delivered ASOs had the smallest global effect on the bacterial transcriptome (30). Third, it was recently shown that upon translocation into bacteria, the peptide is cleaved within the periplasm leaving the ASO or truncated peptide-ASO conjugates for further uptake into the cytoplasm (29). This minimizes effects of the carrier peptide on target binding.

The conjugates were produced by two different methods: PNA by whole-peptide synthesis (Figure 1B) and all other ASOs by copper-free click chemistry (see Materials & Methods, Figures 1C and S5). Consequently, we included two different delivery peptides as controls in the experiments described below: (KFF)_3_K for PNA and (KFF)_3_KK(N_3_) for all other ASOs.

### RNA-MOE, LNA and PNA have comparable binding affinity to the target mRNA in vitro

To compare the ability of the unconjugated ASOs to hybridize to the target transcript, we used electrophoretic mobility shift assays (EMSAs) and microscale thermophoresis (MST). We first conducted EMSAs using a 40 nucleotide (nt) target RNA sequence (nucleotide −20 to +20 relative to the *acpP* translational start codon), which was 5’-labelled with a Cy5 dye. Sequence-specific ASOs were added at defined ASO:RNA ratios, while water or a scrambled PNA (PNAscr) were used as negative controls. Six of the seven on-target ASOs annealed to the target mRNA, resulting in a clear band shift (Figures 2 and S2). The apparent size of the shifted complexes differs due the molecular weight and charge of the ASOs (Table 2). No difference in hybridization was observed between the two ASO concentrations tested, demonstrating that these ASOs are capable of efficiently binding its target at 1.25x molar excess. In contrast, DNA-PTO and the PNAscr control did not interact with the target mRNA. The DNA-PTO modification is known to reduce the binding affinity of the ASO to its target RNA (55) and it is likely that a 10mer is too short to mediate detectable interactions under our experimental conditions because the DNA-RNA duplex is inherently less stable than the RNA-RNA duplex.

To quantify the binding affinities of the ASOs to the target mRNA, we used MST, a sensitive technique that measures a temperature-induced change in fluorescence of a substrate as a function of the concentration of a non-fluorescent ligand (56,57). While the EMSAs provided a qualitative measurement of target binding of the different ASOs, MST allowed us to quantify target affinity by calculating EC_50_ values, the half maximal effective concentration of binding to target mRNA (Table 2, Supplementary Figure S3). RNA-MOE and LNA showed the highest target affinity with an EC_50_ in the upper picomolar range, followed closely by PNA. In contrast, the binding affinities of RNA-OMe and PMO were ten times lower, with EC_50_ values in the lower nanomolar range (Table 2). Thus, RNA-MOE and LNA bound to target mRNA at affinities comparable to the neutral PNA. Both are known to adopt C3‘-endo ribose pucker conformation, which enhances their binding affinity to target mRNA (58,59).

### RNA-MOE, LNA and PNA are strong inhibitors of translation in a cell free system

As a next step, we used a cell-free *in vitro* translation system (31) to investigate whether ASO binding to mRNA leads to a block in protein synthesis. As template, we used the *acpP* sequence spanning −40 to + 51 relative to the AUG start codon fused to a *gfp* reporter gene (*acpP*::*gfp*). This construct was *in vitro* transcribed, pre-incubated with different ASOs at different ratios and subsequently translated *in vitro*. We measured the abundance of the translated AcpP(1–17)-GFP fusion protein by western blot. Water and PNAscr were used as negative controls. In line with their high binding affinities, RNA-MOE, LNA and PNA were effective inhibitors, substantially blocking *in vitro* translation of *acpP*::*gfp* at 1.25x molar excess (Figures 3 and S4). Inhibition of translation *in vitro* was less pronounced in the presence of RNA-F, PMO or RNA-OMe. As expected, DNA-PTO, which is unable to bind to the target mRNA under our experimental conditions, did not inhibit protein synthesis (Figure S4).

**Figure 3.**
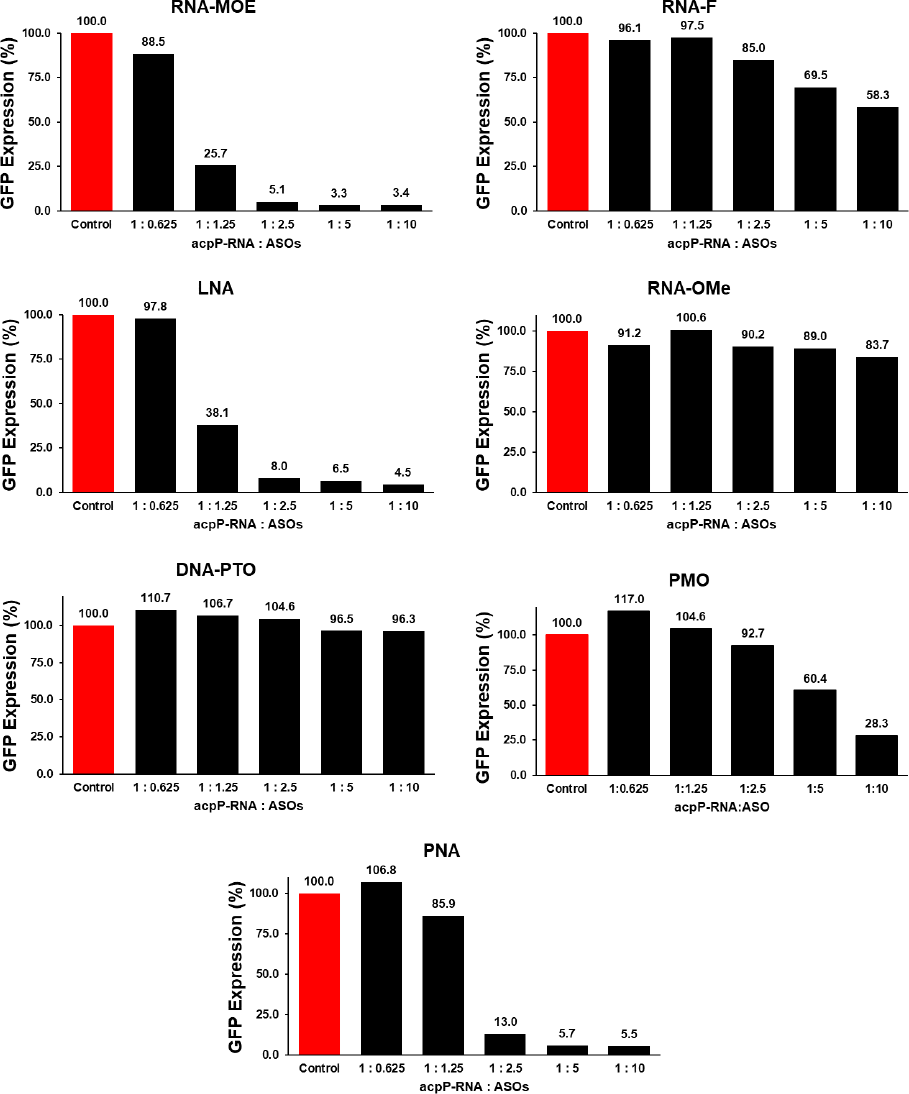
*In vitro* translation of target mRNA. The capacity of the ASOs to inhibit *in vitro* translation of an *acpP::gfp* reporter transcript was analyzed in 10:1, 5:1, 2.5 :1, 1.25:1 to 0.6:1 molar ratios of ASO:RNA. Water served as negative control (Control). Graphs show Image J quantitation of the GFP fusion protein in a western blot analysis. Protein expression levels are shown relative to the water control. The corresponding western blots are shown in Figure S4.

**Figure 4.**
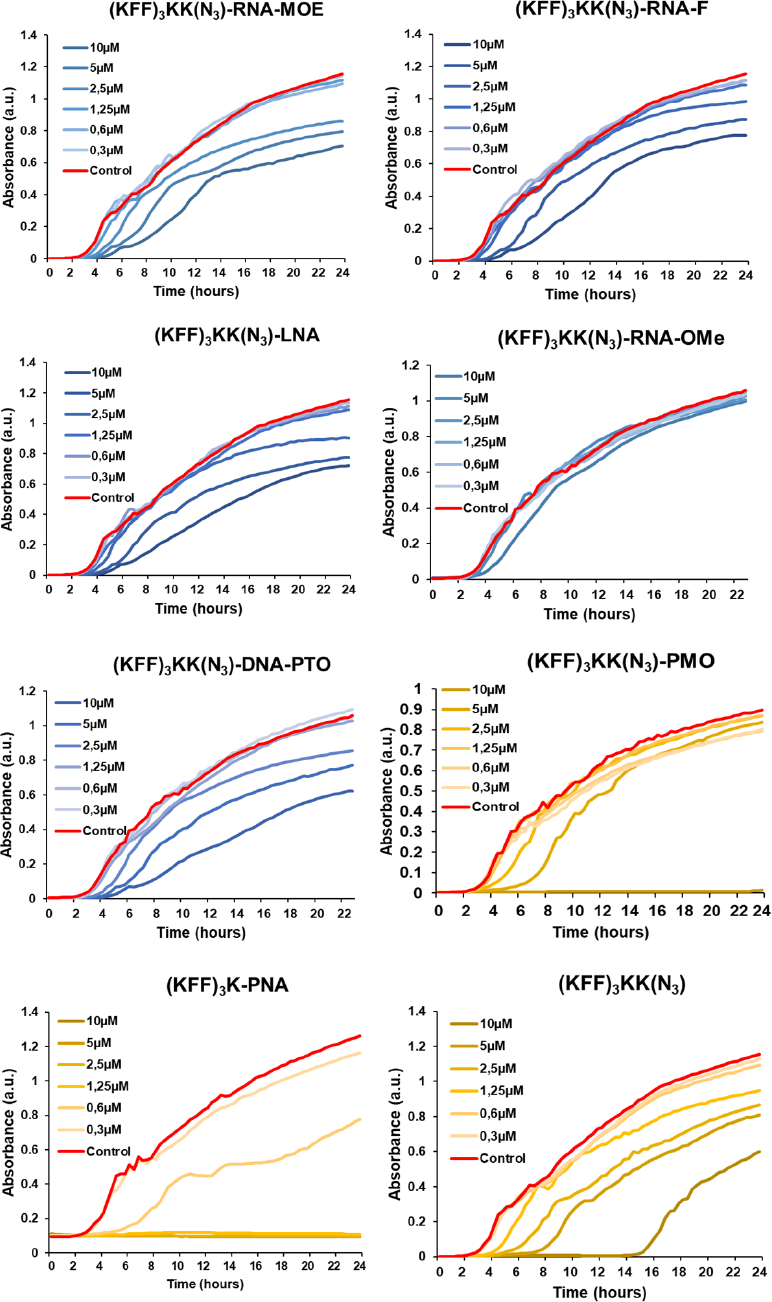
Antibacterial activity of KFF-ASO conjugates. Growth of *Salmonella* was monitored in presence of varying concentrations of KFF-ASO conjugates. The colour coding indicates neutral backbones or the cationic carrier peptide controls (shades of yellow) or anionic phosphate backbones (shades of blue). Water control is shown in red. Graphs are representative examples of two independent experiments.

### Only PNA and PMO show antibacterial activity

Next, we sought to compare the antibacterial efficacy of the ASOs upon delivery into bacterial cells with a KFF carrier peptide. Therefore, we measured the growth kinetics of *Salmonella* in the presence of the ASOs for 24 hours (Figure 4). It is known that the KFF peptide starts showing antibacterial activity at high micromolar concentrations (60). In order to negate the effect of the peptide on the antibacterial efficacy, we considered 10 µM as a cut-off point in our experiments. As expected, none of the unconjugated ASOs showed any antibacterial activity against *Salmonella* (Figure S6), reiterating the importance of the carrier peptide. When delivered by the KFF peptide, PNA inhibited the growth of *Salmonella* at 1.25 µM (as reported before (30)) and PMO did so at 10 µM (Figure 4). However, none of negASOs completely inhibited bacterial growth at the concentrations tested, although a slight growth retardation was observed at 5 µM and 10 µM for RNA-MOE, LNA, RNA-F and DNA-PTO conjugates. The RNA-OMe conjugate showed no activity at all. The fact that the DNA-PTO conjugate, which failed to bind and inhibit the *acpP* target *in vitro*, did show growth retardation at 5 µM and 10 µM is likely due to contributions of the KFF peptide moiety. Indeed, the unconjugated peptide also inhibited *Salmonella* growth at 10 µM until 12 h. In conclusion, RNA-MOE and LNA do not effectively inhibit bacterial growth despite their high binding affinities for target mRNA in cell-free systems, while PNA and PMO do.

### Global transcriptomic changes upon treatment of Salmonella with different ASO modalities

To assay bacterial responses to the different ASO modalities and to determine target mRNA depletion *in cellulo*, we performed RNA-seq analysis of *Salmonella* challenged with 5 µM KFF-ASO conjugates for 15 min (Figures 5-8). We first evaluated two individual RNA-seq experiments by principal component analysis (PCA) and observed three distinct clusters (Figure 5).

**Figure 5.**
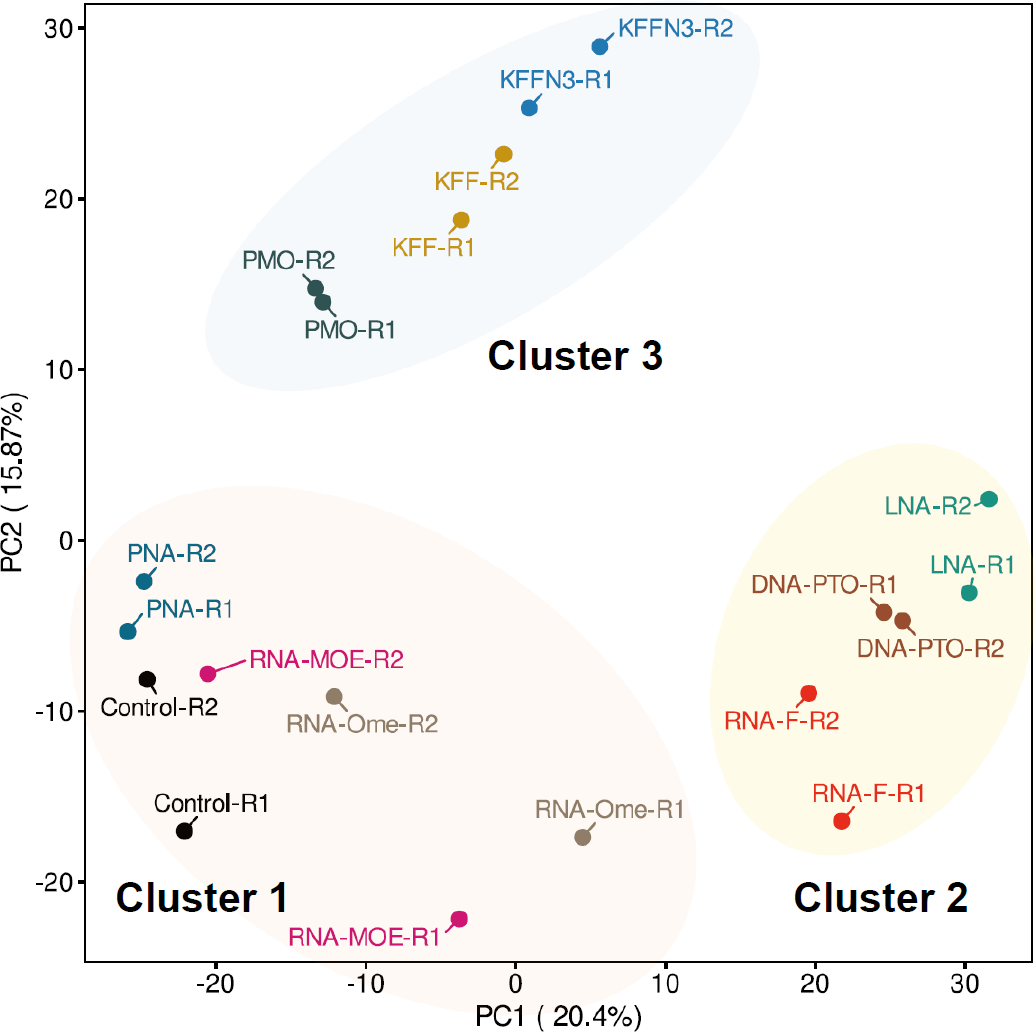
Trascriptomic analysis of KFF-ASO treated *Salmonella*. Principal component analysis (PCA) of two independent RNA-Seq experiments. The PCA plot shows a projection of the RNA-seq data onto the first two principal components. Treatment conditions separate into three clusters, as shown by manual addition of cluster-ellipses in the PCA plot.

**Figure 6.**
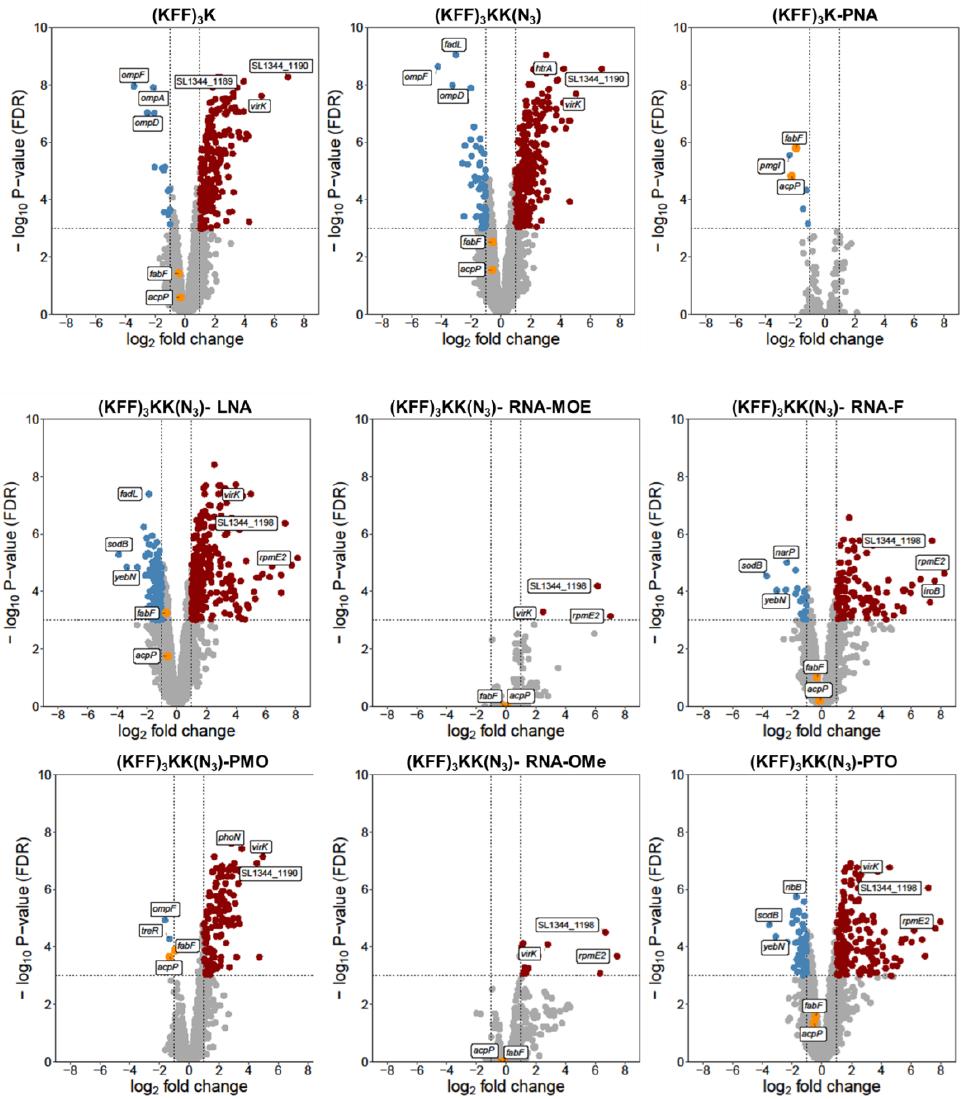
Transcriptomic responses of *Salmonella* upon KFF-ASO treatment. Volcano plots show calculated changes in *Salmonella* gene expression as false discovery rate (FDR)-adjusted *p*-value (-log_10_, y-axis) and fold change (log_2_, x-axis). Significantly differentially regulated genes are characterized by an absolute fold change >2 (down-regulated log_2_ < −1, up-regulated log_2_ > 1; vertical dashed line) and an FDR-adjusted *p*-value < 0.001 (-log_10_ > 3, horizontal dashed line). Significantly down-regulated genes are highlighted in blue, up-regulated genes are highlighted in red. The top-3 up- or down-regulated transcripts are labelled.

**Figure 7.**
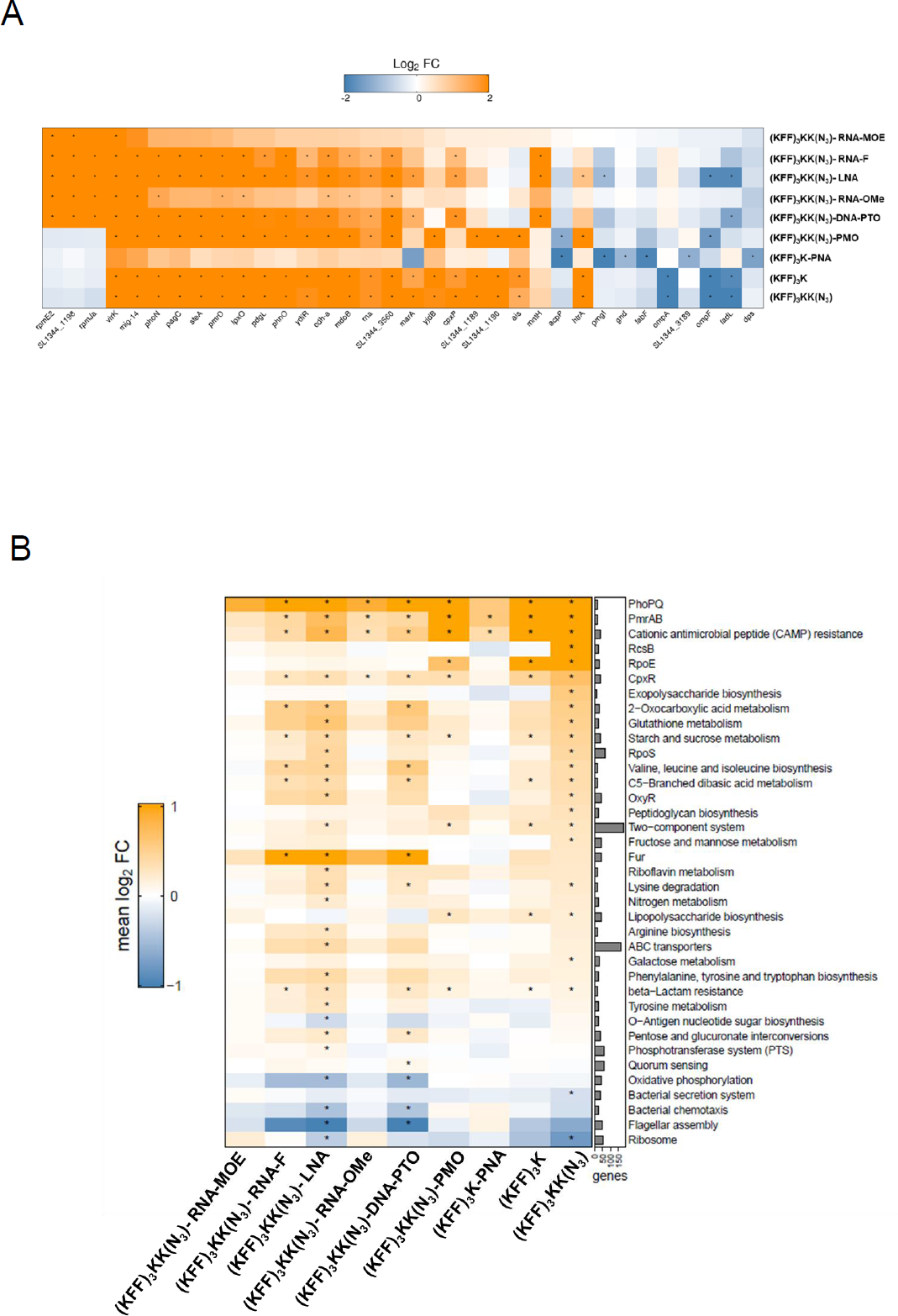
Analysis of differentially expressed genes, KEGG pathways and regulons. (A) Heatmap showing the most strongly differentially expressed genes for all KFF-ASO conjugates. Duplicate RNA-seq samples of *Salmonella* treated for 15 min with KFF-ASOs were normalized to untreated control samples. Log_2_ fold changes (FC) are indicated by color. Orange and blue colors indicate up- and downregulation, respectively. The heatmap includes the top 10 regulated transcripts per condition. Asterisks (*) denote significantly regulated (absolute log_2_FC >1 and false discovery rate (FDR) adjusted P-value <0.001) genes in the respective condition. (B) The heatmap shows the mean log_2_FC for genes assigned to the gene sets. Only the 10 most significantly enriched/depleted gene sets per condition are visualized. Asterisks (*) denote statistical significance (FDR adjusted P-value <0.01) in the respective condition. The bar chart on the right shows the number of genes assigned to the respective gene set.

**Figure 8.**
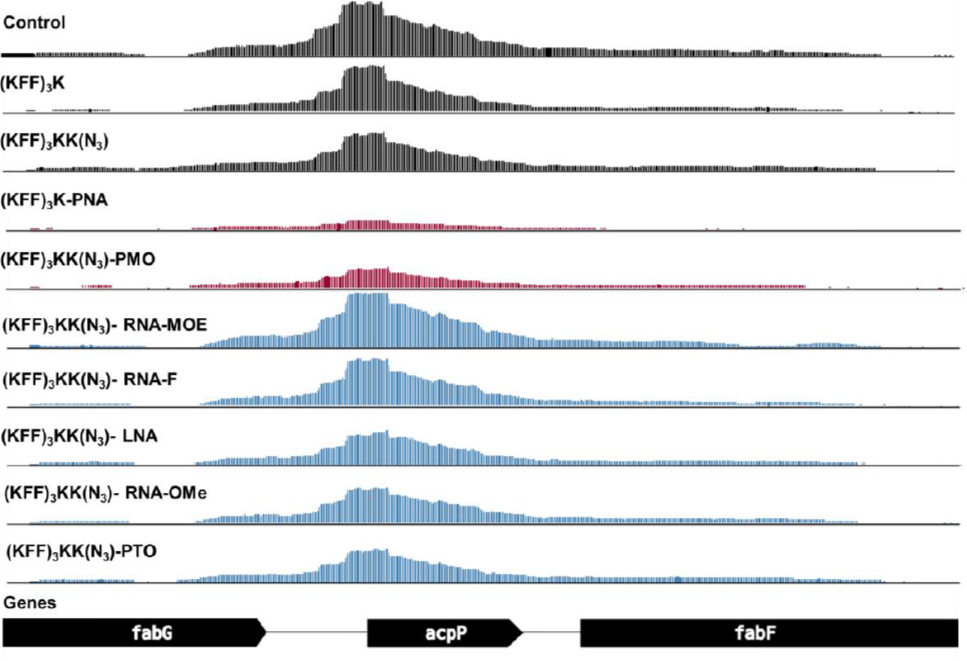
Coverage plot of the *acpP* transcript and neighboring genes for all tested RNA-seq conditions. The coverage plot shows the abundance of mapped reads normalized by counts per million (CPM). The y-axis of all tracks shows the normalized read depth per position, ranging from 0 to 1,500 CPM. Control samples without ASO, ASO modalities triggering significant (log2FC < −1 and FDR < 0.001) *acpP* depletion and other ASO modalities are colored in black, red, and blue, respectively. One replicate (R1) per sample of the RNA-seq data is shown.

The PCA cluster 1 grouped the water control, RNA-MOE, RNA-OMe, and the PNA treated conditions. The treatments in this cluster lead to only small changes in gene expression as compared to the negative control (Figure 5), confirming our previous observation that KFF coupled to PNA has little effect on the global transcriptome (30), and indicating that RNA-MOE and RNA-OMe do not trigger strong transcriptomic responses either (Figure 6). Cluster 2 grouped the other negASOs. As we have previously observed for KFF peptide exposure (30,31), these treatments led to activation of a cationic antimicrobial peptide (CAMP) response, including induction of the PhoP/Q and PmrA/B regulons involved in remodelling LPS, and the CpxR regulon that responds to envelope stress (Figure 7). This indicates that these compounds interact with the cell membrane. These treatments also led to downregulation of genes responsible for destruction of superoxide anion radicals (*sodB)* and those involved in manganese efflux *(yebN)* (Figures 6 and 7). Finally, cluster 3 grouped samples of the unconjugated peptides and the PMO treatment. Similarly to cluster 2, these activated a strong CAMP response (Figure 7). The observation of differential activation of membrane stress responses indicates that the cellular response is dependent on both the carrier peptide and the ASO modality and suggests the need for joint optimization of both components in designing ASOs for targeted gene silencing.

Next, we evaluated target mRNA depletion. In agreement with our previous work (30), we found that PNA treatment reduced the levels of *acpP* mRNA (Figures 6-8). This is also true for PMO treatment (Figures 6-8). In contrast, none of the negASOs caused significant depletion of the target transcript (FDR adjusted P-value <0.01), although LNA had a slight effect (Figures 6 and 8). Reduction of *fabF* (3-oxoacyl-[acyl carrier protein] synthase 2), which is co-transcribed with *acpP*, was also observed for PNA and PMO (Figures 6); an effect we have seen before for PNA (30). Therefore, we conclude that only the ASOs that showed antibacterial activity against *Salmonella* induce target mRNA depletion *in vivo* within 15 min of treatment.

### Treatment with PNA and PMO lead to lower AcpP protein expression in Salmonella

We have previously shown that PNAs targeting *acpP* cause depletion of the target transcript. However, a reduction in target protein levels is not always associated with a detectable depletion of the mRNA for all target transcripts (31). Therefore, we aimed to test if the different ASO modalities affect AcpP protein levels *in cellulo* despite their inability to deplete the *acpP* transcript. Thus, we used *Salmonella* expressing FLAG-tagged AcpP (AcpP::3xFLAG) under its native promoter to study the effect of PNA and PMO on its expression. First we exposed the bacterial cells to PMO and PNA for 30, 60 or 120 min (Figure 9A). PNA mediated depletion in AcpP levels can be observed from 60 min post treatment onwards and the effect increases at 120 min post treatment (Figure 9A). We also extended the study to 240 min, but at this time point the PNA starts exerting antibacterial activity. Similarly, treatment with PMO led to gradual depletion of AcpP, albeit less pronounced than PNA. We repeated the experiment for all ASOs in parallel at the 120 min time point, but we did not observe a detectable reduction in AcpP protein levels upon negASO treatment (Figure 9B), in line with their inefficacy in inhibiting bacterial growth.

**Figure 9.**
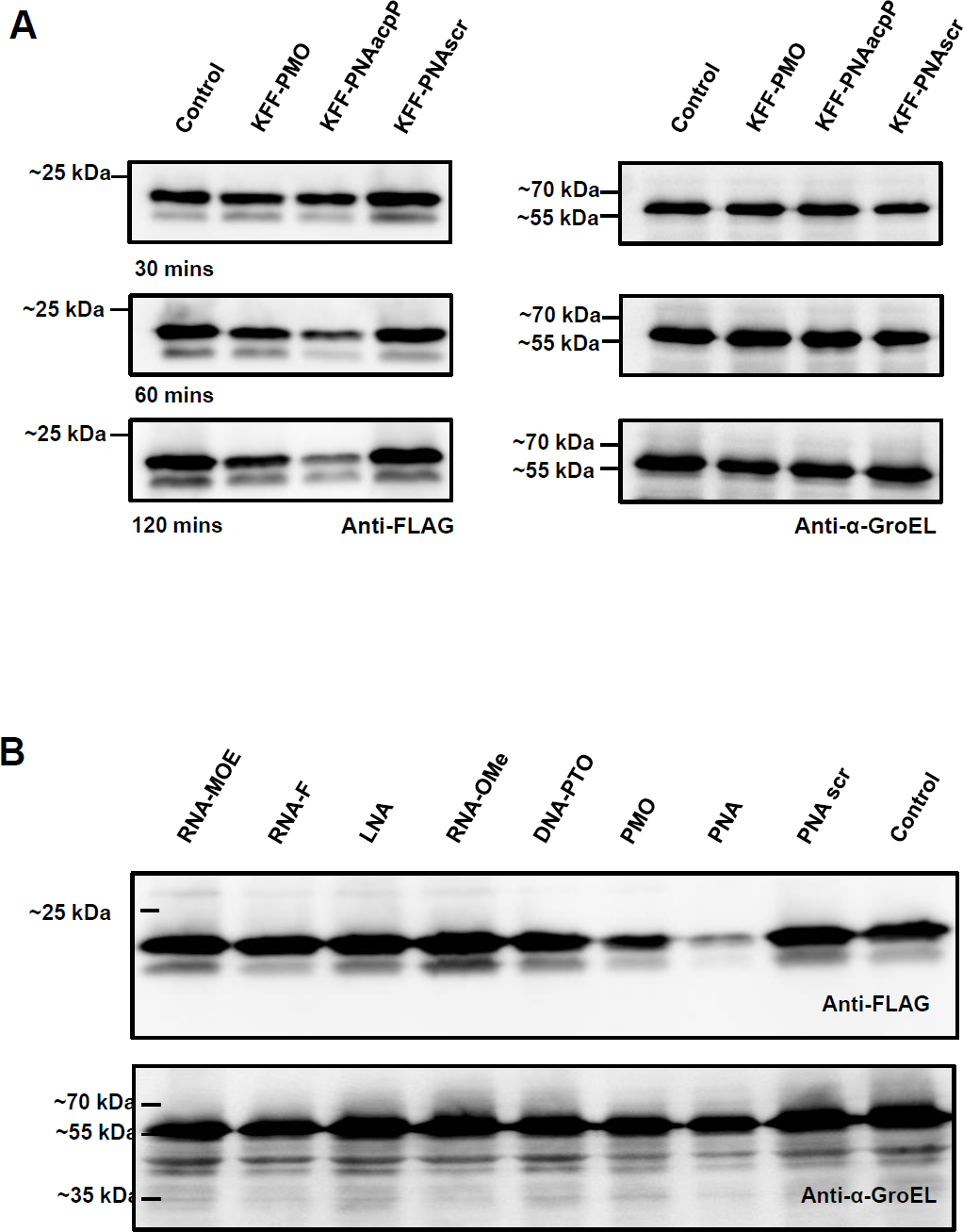
Western blot analysis of AcpP levels upon KFF-ASO treatment of *Salmonella*. (**A**) Western blot time course of KFF-PNA- or KFF-PMO-treated *Salmonella*. (**B**) Western blot analysis of *Salmonella* treated with all KFF-ASO modalities for 120 min. (**A,B**) Blots were probed with a FLAG-specific antibody to detect the AcpP::3xFLAG fusion protein. Mock-treatment (Control) and scrambled PNA (PNA scr) served as controls. GroEL was used as a loading control.

### KFF-mediated permeabilization of the outer membrane of Salmonella is compromised by conjugation with negASOs

Since the negASOs are ineffective in bacterial cells despite their activity in cell-free systems, we investigated if these ASOs affected the ability of the KFF carrier peptide to induce outer membrane (OM) permeabilization. Therefore, we used the hydrophobic fluorophore *N*-phenylnaphthylamine (NPN), a dye that turns fluorescent after binding to hydrophobic regions within cell membranes. This only occurs in cells with a compromised OM (48,50). CTAB (a surfactant) and Polymyxin B (PMXB), known permeabilizers of the bacterial OM, cause a rapid increase in florescence when added to *Salmonella* in this assay (Figure 10). Both KFF control peptides also permeablize the OM, and this ability is retained upon conjugation to PNA and PMO. However, when conjugated to negASOs, the KFF carrier is unable to permeabilize the OM (Figure 10). This is could potentially be due to electrostatic interactions between the negative charged ASO and the positively charged KFF. Overall, these results indicate that conjugation of the negASOs to the KFF peptide compromises its cell membrane permeabilizing activity.

**Figure 10.**
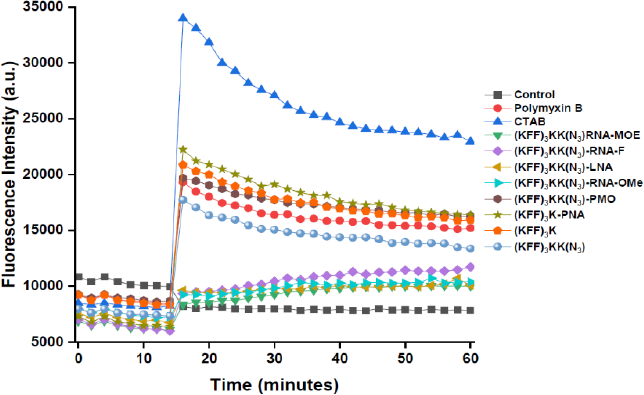
Outer membrane permeabilization of *Salmonella*. NPN fluorescence plotted over time. Different KFF-ASOs and controls were added at 14 min. CTAB and Polymyxin B were used as positive controls, water as a negative control. A representative example of two independent experiments is shown.

### negASO are not delivered efficiently inside Salmonella by the KFF peptide

To test directly if the ASO conjugates translocate into *Salmonella*, we used MALDI mass-spectrometry (MS) to identify ASO fragments in the bacterial periplasm after treatment with 10 µM KFF-ASO conjugates for 15 min (Table 2 and Figures S7-S10). Both PNA and PMO fragments were detected in the periplasmic fraction (Figure S7). The KFF-PNA fragments corresponded to either peptide-cleaved PNA or PNA attached to a single unit of KFF. This is in line with a recent study that reported that the KFF peptide is cleaved within the periplasm prior to translocation into the cytoplasm (29). In the case of PMO, only the PMO fragment was observed in the periplasm, suggestive of KFF cleavage. Importantly, none of the negASOs were detected in these subcellular fractions (Figures S8 and S9), likely explaining the lack of antibacterial effects against *Salmonella*.

## DISCUSSION

Despite their successful clinical application for treating genetic disorders in humans, ASOs are yet to be established as antibacterial drugs. For their effective clinical use, several important questions need to be answered. These include the identification of the best ASO modality, the best ASO carrier, the range of potential target genes and the overall bacterial response to ASO modalities. We have been trying to address these questions systematically with the ultimate goal of developing ASOs as programmable, species-specific antibiotics in complex bacterial communities (1,31,51). In the current study, we focused on a direct comparison of the antisense properties of seven different ASO modalities in the model pathogen *Salmonella*. Our results are provided as an overview in Table 3 and we will discuss our findings and directions for future studies below.

**Table 3.** Summary of the comparison between the different ASOs.

| ASO modality | Binding affinity<br>(*in vitro*) | Inhibition of translation<br>*in vitro* | Antibacterial activity<sup>§</sup> | *acpP* depletion<br>(*RNA-seq*)<sup>§</sup> | AcpP protein reduction | Outer-membrane permeation<sup>§</sup> | Uptake inside bacteria (MS)<sup>§</sup> |
| --- | --- | --- | --- | --- | --- | --- | --- |
| RNA-F | + | ++ | - | - | - | - | - |
| RNA-OMe | ++ | + | - | - | - | - | - |
| RNA-MOE | +++ | +++ | - | - | - | - | - |
| LNA | +++ | +++ | - | - | - | - | - |
| DNA-PTO | - | - | - | - | - | - | - |
| PMO | ++ | ++ | + | + | + | ++ | + |
| PNA | +++ | +++ | ++ | ++ | ++ | ++ | + |

<sup>§</sup> Indicates that the ASO was conjugated to the carrier peptide. Effect strength is represented by additional (+) signs.

### ASO chemistry and binding affinity

Several ASO drugs based on diverse nucleotide modalities including DNA-PTO, RNA-MOE, RNA-OMe, RNA-F and PMO have been approved for the treatment of human diseases (22,61-63). In terms of pharmacokinetics, pharmacodynamics and bioavailability in the human body each modality has different properties (64). In addition, the RNA binding affinity of the ASO is also affected by these modifications. For example, RNAs containing 2’-modified sugars form more stable duplexes, while modifications on the phosphate backbone generally lower duplex stability (65). Here, we compared these different ASO modalities with PNA, currently the most commonly used modality in antimicrobial research. In general, our results are in agreement with the trends observed in structure–stability studies with different chemically-modified nucleotide duplexes (65). With the exception of DNA-PTO, all ASOs annealed to the target mRNA *in vitro*. Phosphorothioate linkages are known to reduce affinity for complementary RNA (57) and it is likely that the DNA-PTO used in our experimental setting is not long enough to mediate efficient binding. LNA and RNA-MOE showed a stronger affinity to target mRNA compared to the neutral PNA and PMO. Therefore, for a length of ten bases, charge contributions from the phosphate backbone do not seem to strongly affect target binding.

### The net charge of KFF-ASO conjugates affect antibacterial activity

Unlike mammalian cell membranes, which are zwitterionic in nature, bacterial membranes have an overall negative charge (66). Therefore, cationic compounds have greater affinity for the bacterial membrane in comparison to anionic compounds (66). The KFF peptide contains four units of the cationic amino acid lysine, and six units of the hydrophobic amino acid phenylalanine. Conjugation of KFF to the charge-neutral PNA or PMO retains the positive charge of the KFF peptide. In contrast, the negASO-KFF conjugates have a net negative charge, potentially hindering their interaction with the bacterial membrane. In addition, electrostatic interactions between the negASO and the KFF peptide can lead to intra- and intermolecular aggregates. This might be the reason why conjugation of CPPs to charged-backbone ASOs has not been widely explored, although there are successful examples of such conjugates (23). Nevertheless, our observation that the negASOs are unable to cross the bacterial OM of *Salmonella* (within 15 min post treatment) when conjugated to KFF indicates that these ASOs might have a negative effect on the membrane-permeating properties of KFF. While some of the negASOs cause growth retardation of *Salmonella* in our experimental settings, this effect cannot be explained by specific reduction of its cellular target on RNA or protein level. However, our RNA-seq approach showed that with the exception of RNA-MOE and RNA-OMe, negASOs trigger the activation of stress response pathways related to PhoP/Q, PMR A/B and CAMP. This indicates that these negASOs have some interaction with the cell membrane and might lead to stress-induced toxicity.

### negASOs require alternative delivery vehicles

Our data show that LNA and RNA-MOE are potent translational inhibitors of the *acpP* target transcript *in vitro*. In fact, they outperform PNA and PMO in this assay. However, they are not efficiently delivered across the bacterial membrane by the KFF carrier peptide. Replacement of the KFF peptide with peptides that bear more cationic amino acids might be more effective in delivering negASOs. Alternatively, the negative charges of the phosphate backbone can also be used to complex negASOs with other cationic carriers such as bolaamphiphiles (19). This will mitigate the net negative charge of the complex and may lead to better delivery. There have been reports of successful delivery of ASOs across the bacterial membrane with vitamin B_12_, DNA nanocages, bolaamphiphiles and lipid nanoparticles (27,61,67-70) and recent advances in the design of macromolecular carriers might open up more avenues for the use of negASOs as antibacterials. These are currently rarely applied in the field of antisense antibiotics, but the results described in this manuscript argue for a search for alternative delivery systems beyond CPPs.

### PNA versus PMO

Adopting KFF as a carrier, only the charge-neutral ASOs exhibit strong antibacterial activity against *Salmonella*. While PNAs and PMOs have been compared before with respect to synthesis, stability, backbone flexibility, aqueous solubility, sequence specificity and target binding (71), there has not been a direct comparison of their antibacterial activity. In general, PNAs have a higher binding affinity to RNA, while PMOs are more soluble in water (71). Here, we find that PNA is more effective than PMO in inhibiting translation of *acpP* both *in vitro* and within *Salmonella*. Of note, the molecular weight of the PMO conjugate is ∼1.4 kDa higher than that of the PNA conjugate. Whether this has any influence on bacterial internalization is unclear.

Another point of consideration is the effect of the linker that was used to conjugate PMO to the KFF peptide in this study. PNA-KFF conjugates can be synthesised as fusion peptides due to the PNA pseudo-peptide backbone. In contrast, the PMO conjugate was linked by copper-free click chemistry in our study. It is therefore structurally different from the peptide-PMO conjugates reported in the literature where the peptide is conjugated to the PMO using amide-coupling chemistry (72,73). While there is no direct comparison of the activity between click-coupled and amide-coupled peptide-PMO conjugates, we do not observe a large difference in antibacterial activity of click-coupled PMO compared to an earlier study using amide-coupled PMO (72). Thus, the linker is unlikely to strongly affect the antimicrobial activity of PMO.

Interestingly, the transcriptomic response of *Salmonella* to PMO is different compared to PNA. While PNA has little effect on the global transcriptome at least when coupled to the KFF peptide, PMO activates membrane stress responses, similar to the unconjugated peptide controls. In addition, PNA has a stronger effect than PMO on the transcript levels of *acpP*, which is also reflected on the protein level. Therefore, in our experimental setting using *acpP* as a target and treating bacteria in culture broth, PNA outperforms PMO as an antisense antibacterial. However, it should be stressed that PMO has shown great efficacy in animal experiments (9,11), which warrants a more systematic comparison of PNA versus PMO with a larger number of targets in the future.

### Limitations of the study

Here, we have compared the antibacterial potential of different ASOs *in vitro* and in bacterial cell culture, using a single target mRNA and a fixed number of bases, limiting conclusion with respect to other targets and ASO sequence lengths. Our study also does not address the pharmacodynamics and pharmacokinetics of the peptide-ASO conjugates within the body of an animal. It would therefore be interesting to compare the effectiveness of the different ASO modalities in animal models of infection. In addition, we acknowledge that a change in peptide carrier might lead to an improvement in antibacterial of some of the negASOs. Indeed, our observations should direct future work towards development of alternative carriers that will enable effective delivery of the negASOs, especially RNA-MOE and LNA.

## DATA AVAILABILITY

Our RNA-seq data set has been deposited with GEO under accession number GSE232819 (https://www.ncbi.nlm.nih.gov/geo/query/acc.cgi?acc=GSE232819). The code used for data analysis is freely available at https://github.com/BarquistLab/aso_screen_ghosh_et_al_2023.

## SUPPLEMENTARY DATA

**Supplementary Data are available online.**

## ACKNOWLEDGEMENT

We thank Phuong Thao Do for extensive technical assistance and Tobias Kerrinnes for laboratory organization. We especially thank Anke Sparmann for editing the manuscript. Anuja Kibe is gratefully acknowledged for her help towards setting up the MST experiment and fruitful discussions for interpreting the results. Juliane Adelmann for helping us with MALDI mass spectrometry. We thank the Core Unit SysMed at the University of Würzburg for excellent technical support, library preparation and RNA-seq data generation (supported by the IZKF at the University of Würzburg, project Z-6).

## FUNDING

Research was supported by the Bavarian Bayresq.net (L.B., J.V.). Funding for open access charge: Bayresq.net and library of the University of Würzburg.

## CONFLICT OF INTEREST

The authors declare no conflicts of interest.

## Notes

https://www.ncbi.nlm.nih.gov/geo/query/acc.cgi?acc=GSE232819

https://github.com/BarquistLab/aso_screen_ghosh_et_al_2023

### Competing Interest Statement

The authors have declared no competing interest.

