## Supporting information for "A side-by-side comparison of peptide-delivered antisense antibiotics employing different nucleotide mimics"

### Figure S1

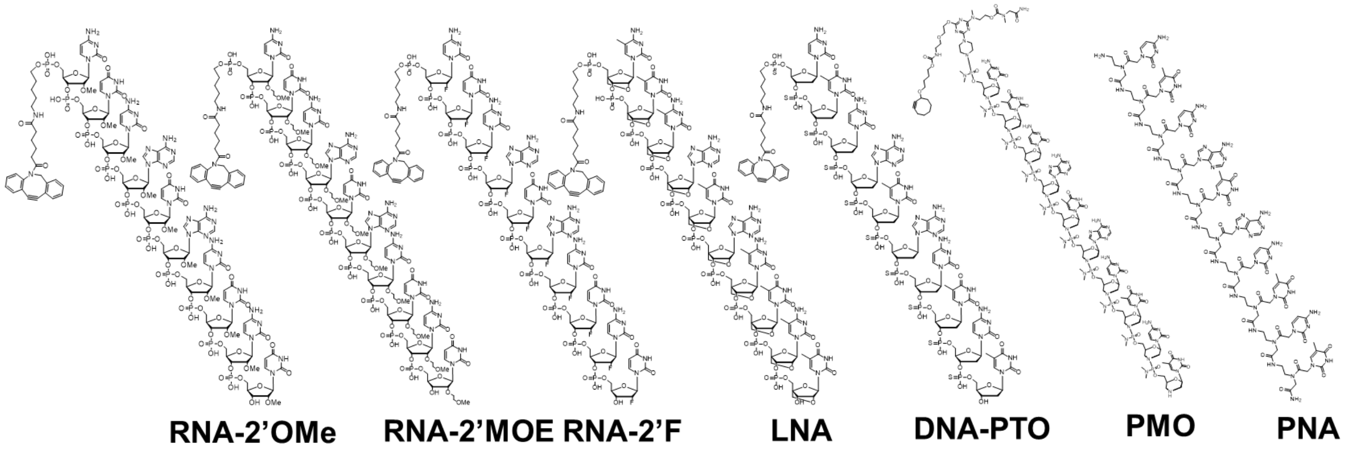


**Supplementary Figure S1**. Chemical structures of ASOs used in the study.

### Figure S2

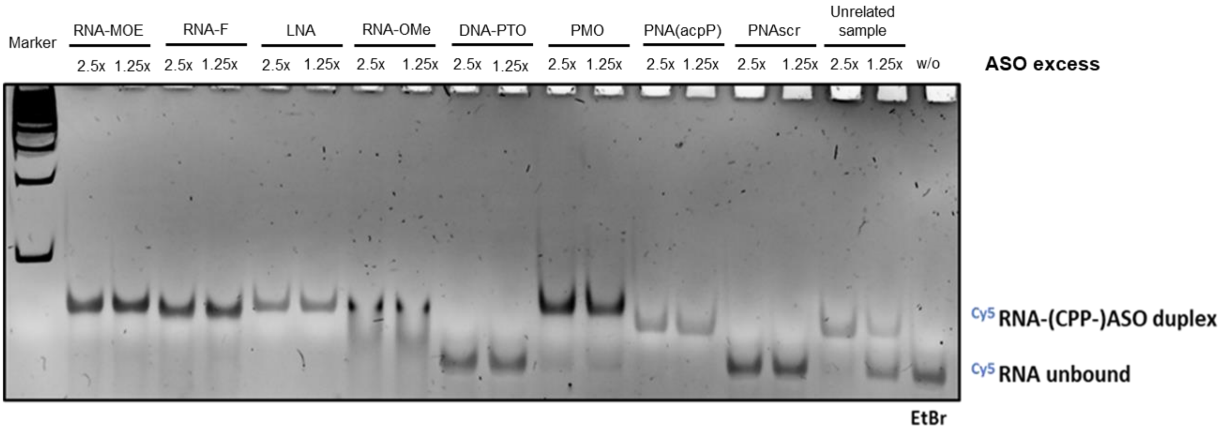


**Supplementary Figure S2.** EtBr staining of the gel presented in Figure 2.

### Figure S3

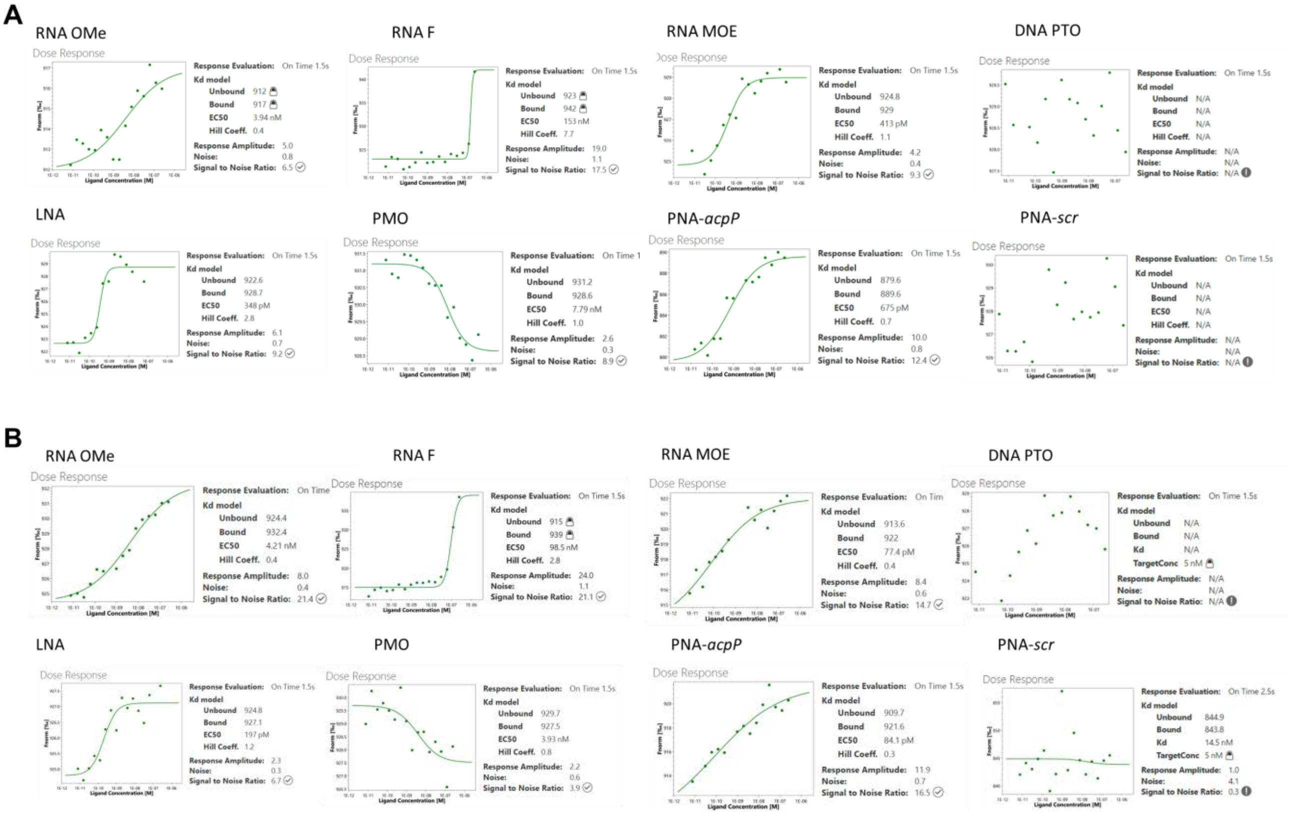


**Supplementary Figure S3**. Microscale thermophoresis assay with all the compounds. PNA with a scrambled sequence was used as a negative control. Data of two independent experiments (presented in panel A and B) were analyzed to obtain ΔFnorm values using the MO Affinity Analysis software (NanoTemper Technologies Gmbh, Munich, Germany). The fitting for $EC_{50}$ values was done with the Hill equation using the MO Affinity Analysis software. The $EC_{50}$ values presented in Table 1 are the mean ± S.D. of two independent experiments. Please note that the opposite trend of the binding curve observed for the PMO sample is in line with the general observations of MST technology. The 'direction' of the curve is dependent on the specific properties of the molecules and can therefore differ between molecule. It does not influence further analysis.

### Figure S4

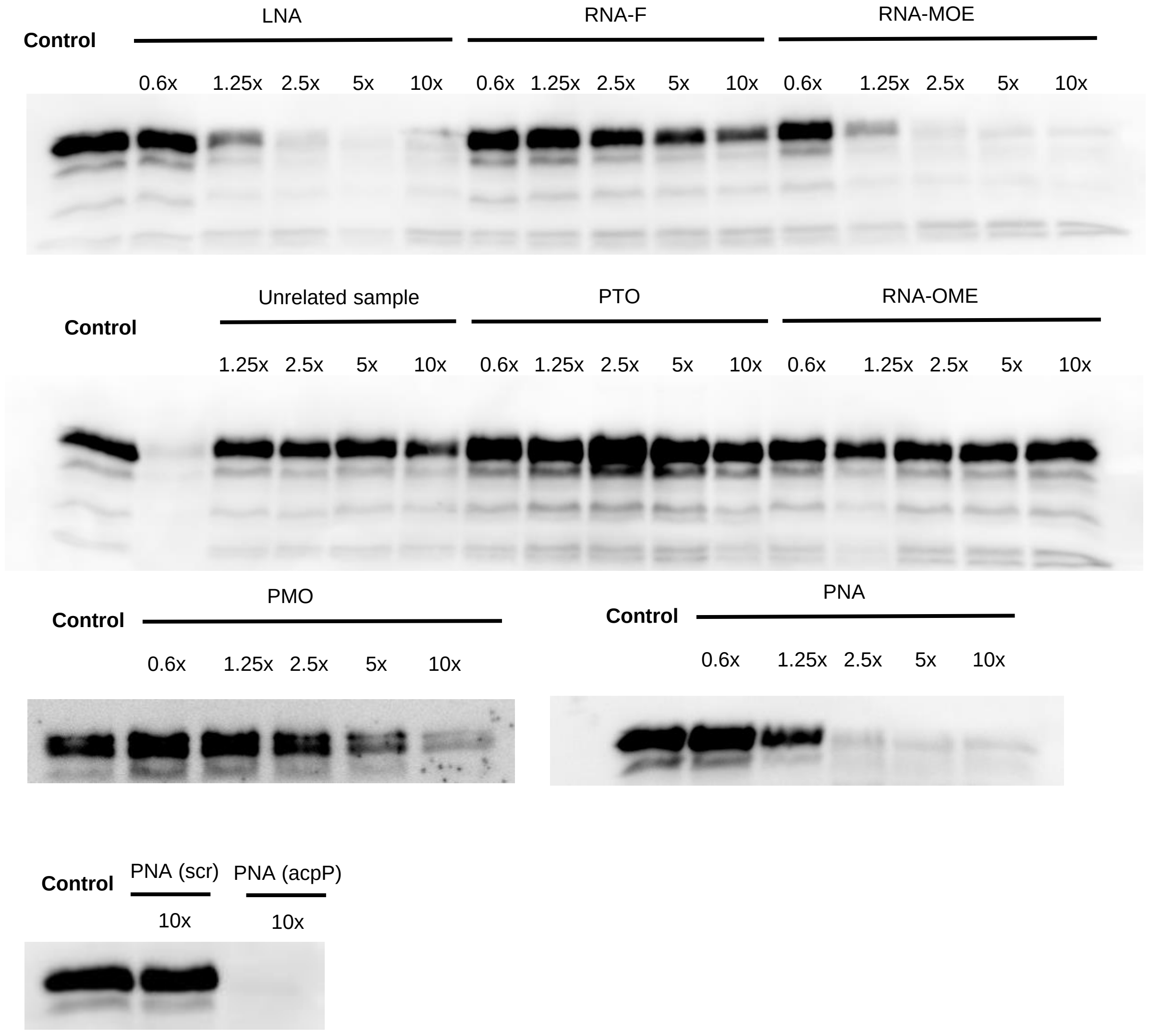


**Supplementary Figure S4**. *In vitro* translation assay of the compounds. Control used here is water.

### Figure S5

A

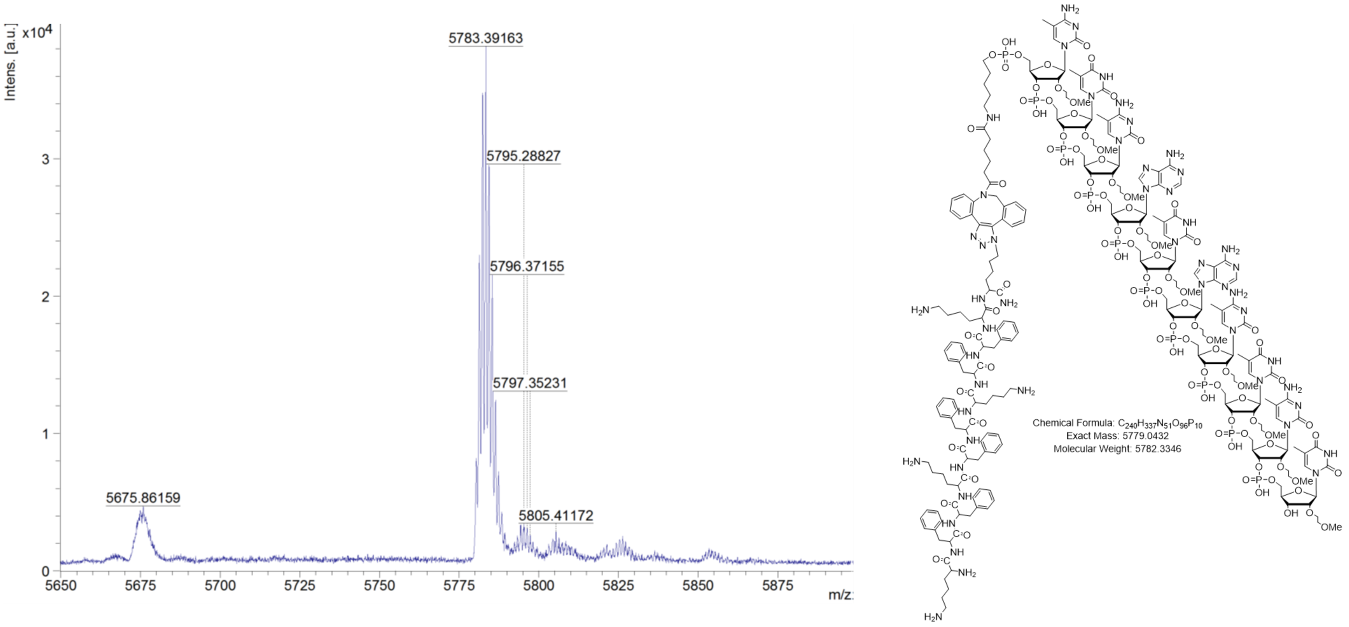

Intens. [a.u.]
x10^4
5783.39163
5795.28827
5796.37155
5797.35231
5675.86159
5805.41172
m/z
Chemical Formula: C240H337N51O96P10
Exact Mass: 5779.0432
Molecular Weight: 5782.3346


B

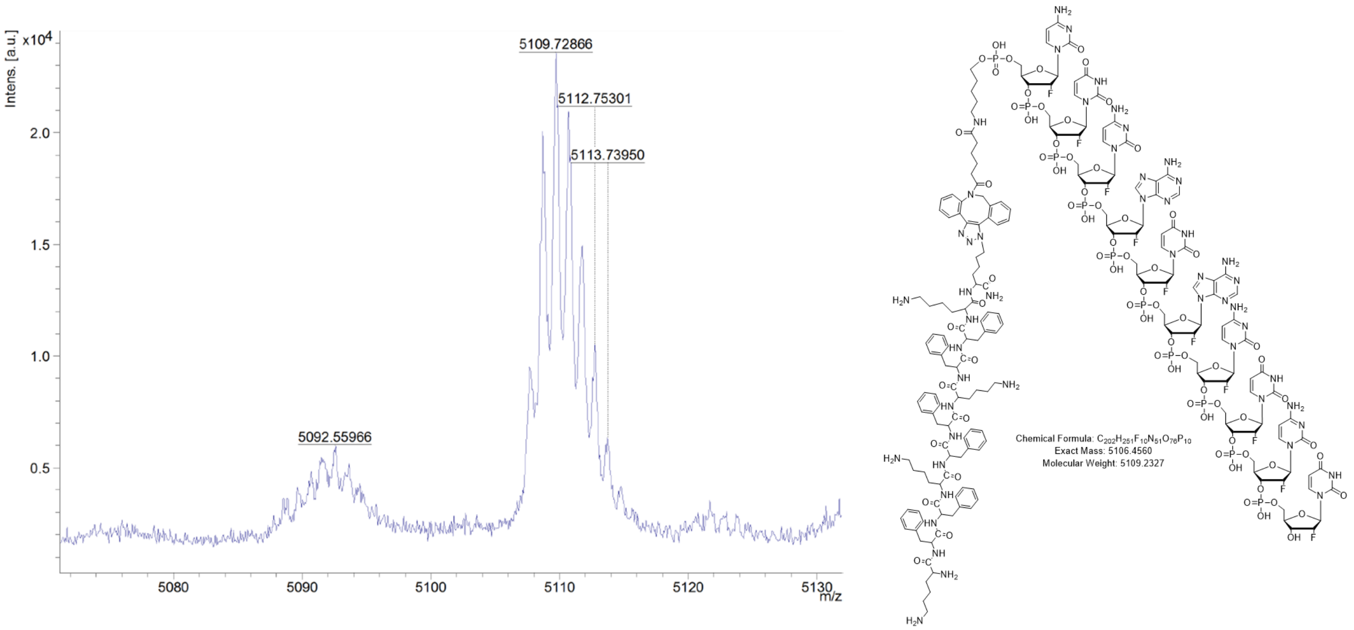

Intens. [a.u.]
x10^4
5109.72866
5112.75301
5113.73950
5092.55966
m/z
Chemical Formula: C202H251F10N51O76P10
Exact Mass: 5106.4560
Molecular Weight: 5109.2327

C

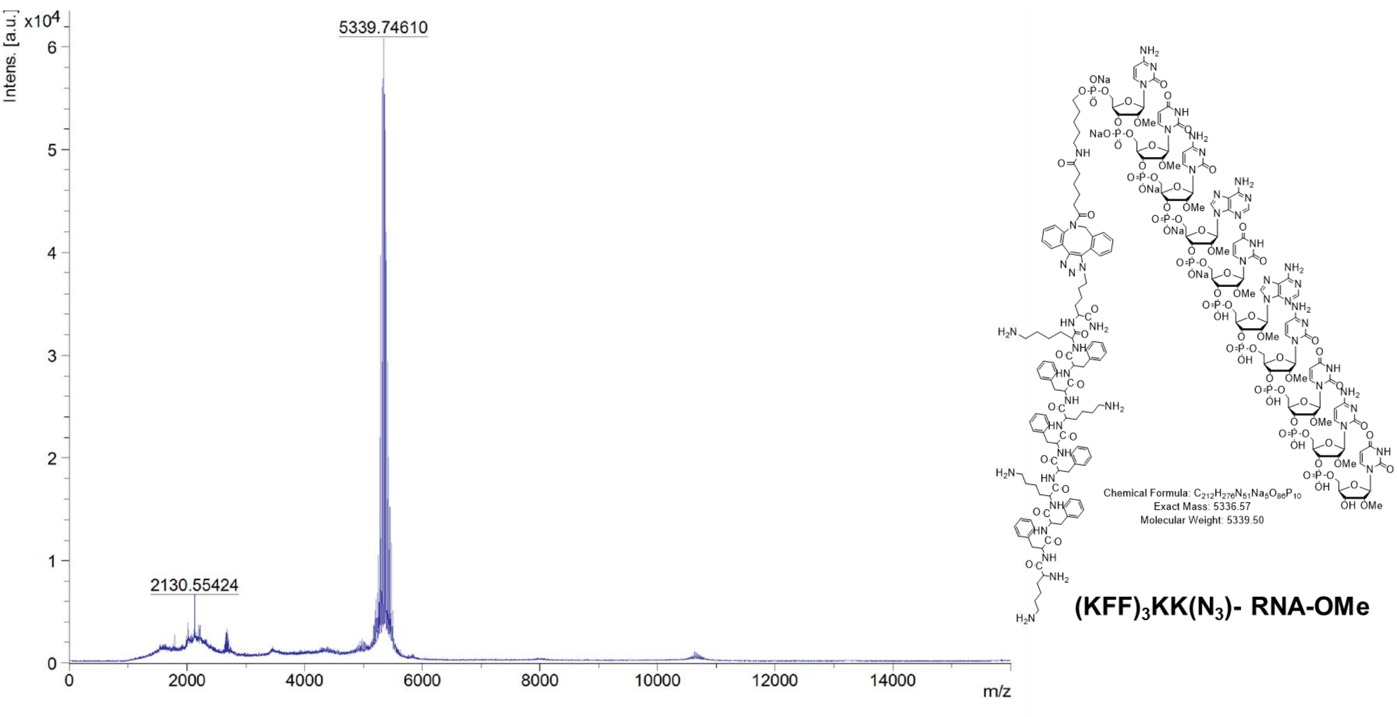
Intens. [a.u.] x10^4
5339.74610
2130.55424
m/z
Chemical Formula: C212H276N51Na5O66P10
Exact Mass: 5336.57
Molecular Weight: 5339.50
(KFF)3KK(N3)- RNA-OMe

D

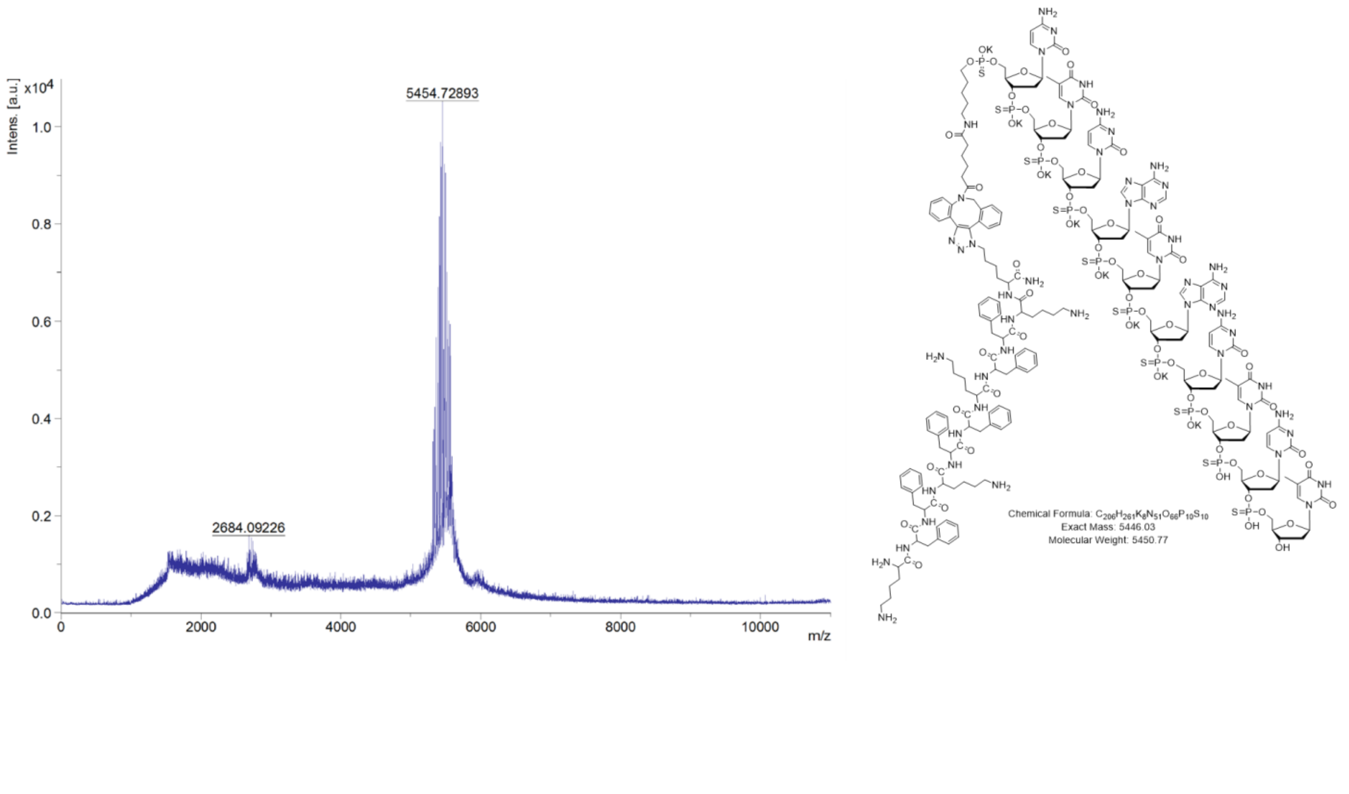
Intens. [a.u.] x10^4
5454.72893
2684.09226
m/z
Chemical Formula: C206H261K8N51O66P10S10
Exact Mass: 5446.03
Molecular Weight: 5450.77

E

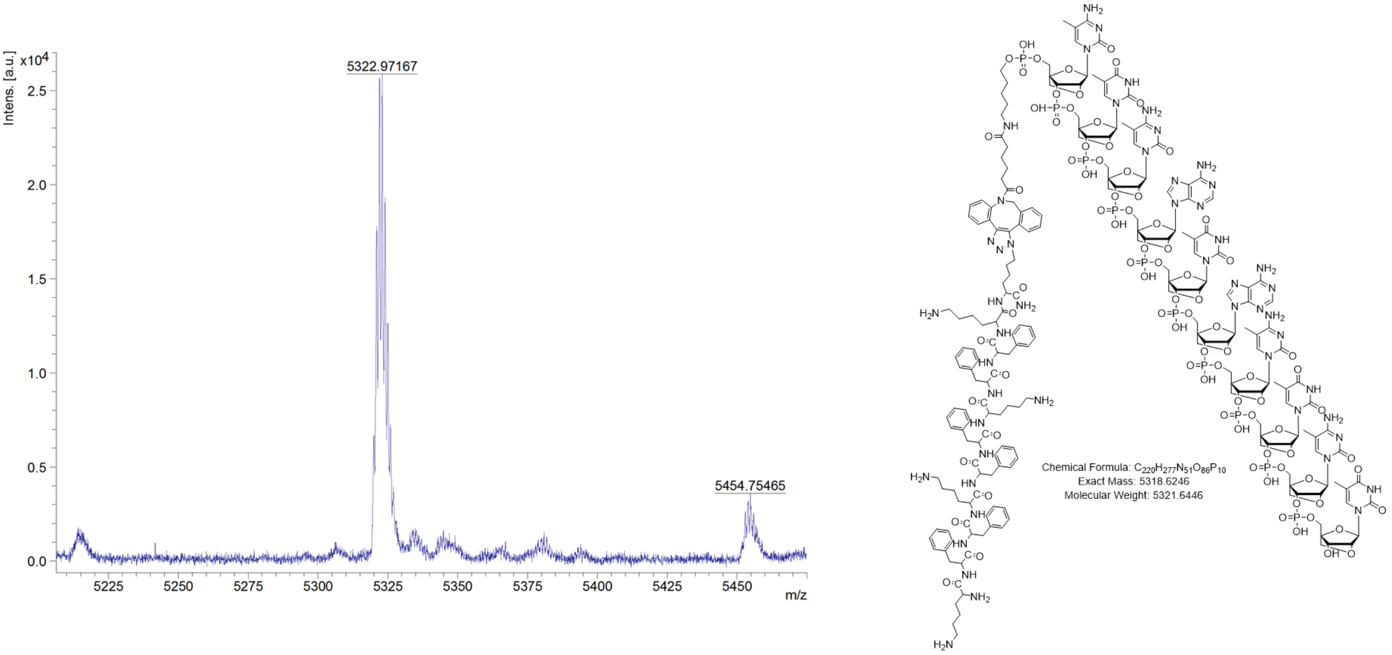


F

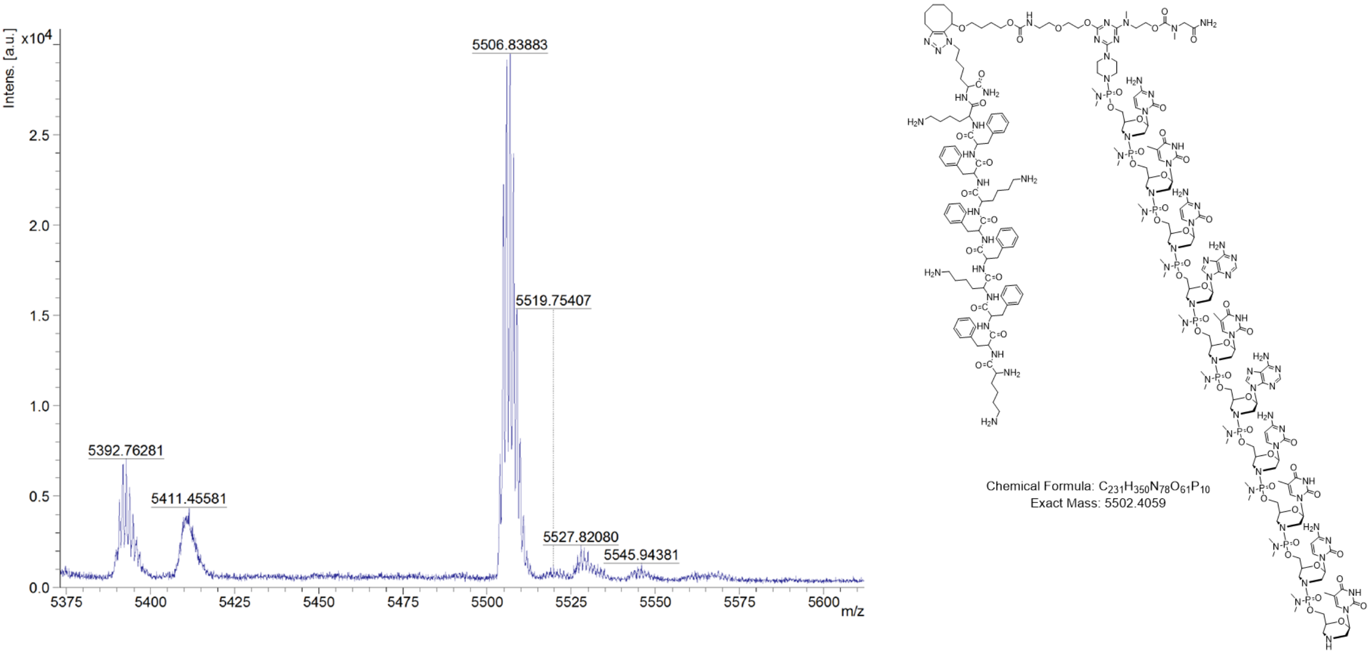


**Supplementary Figure S5.** Structure and MALDI-TOF spectra of conjugates of (KFF)3KK(N3) to RNA-MOE (A), RNA-F (B), RNA-OMe (C), PTO (D), LNA (E) and PMO (F).

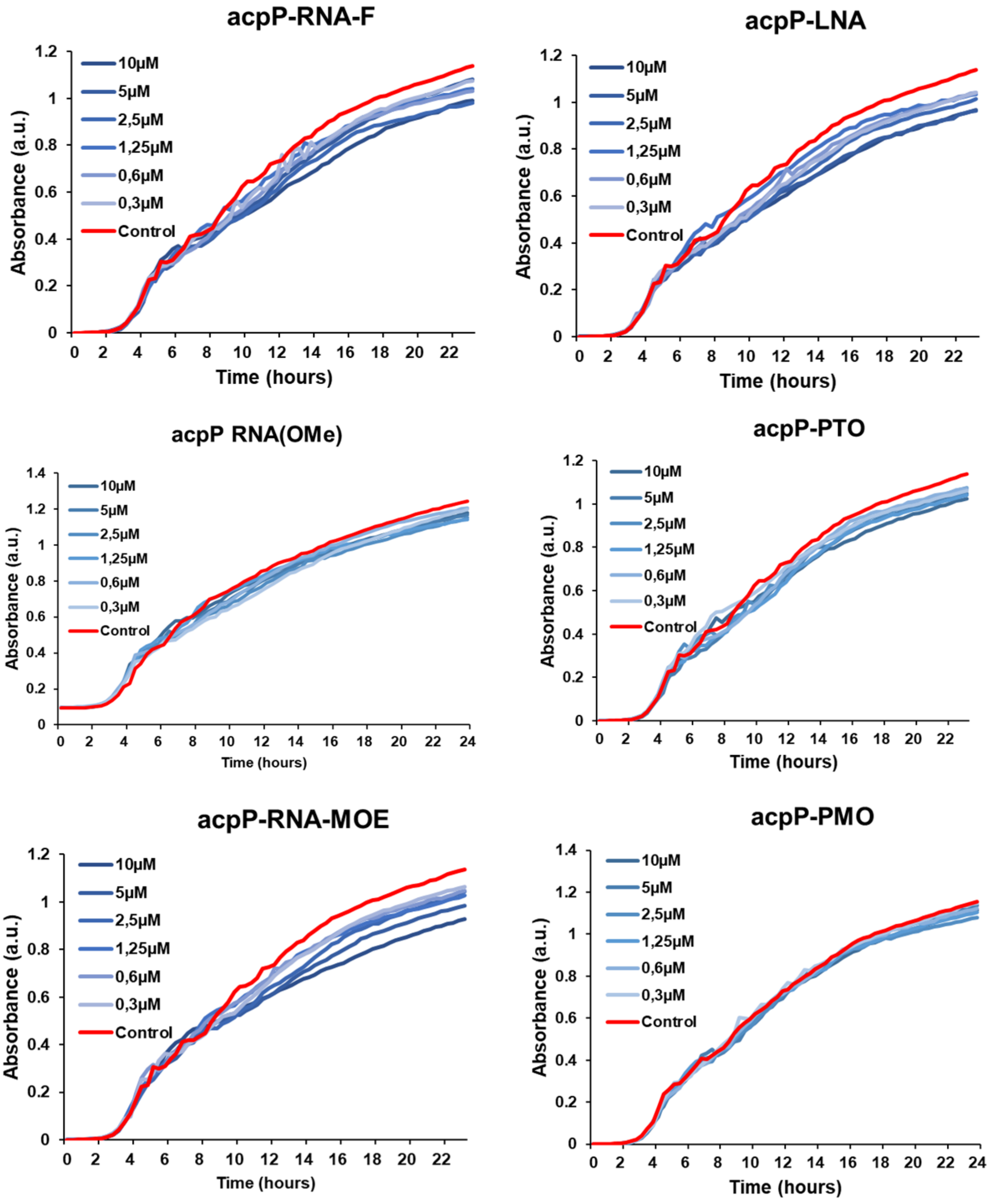


**Supplementary Figure S6**. Growth kinetics of *Salmonella* treated with unconjugated ASOs. Unconjugated PNA is not active against Salmonella (see *Nucleic Acids Res,* 2021, 49, 4705).

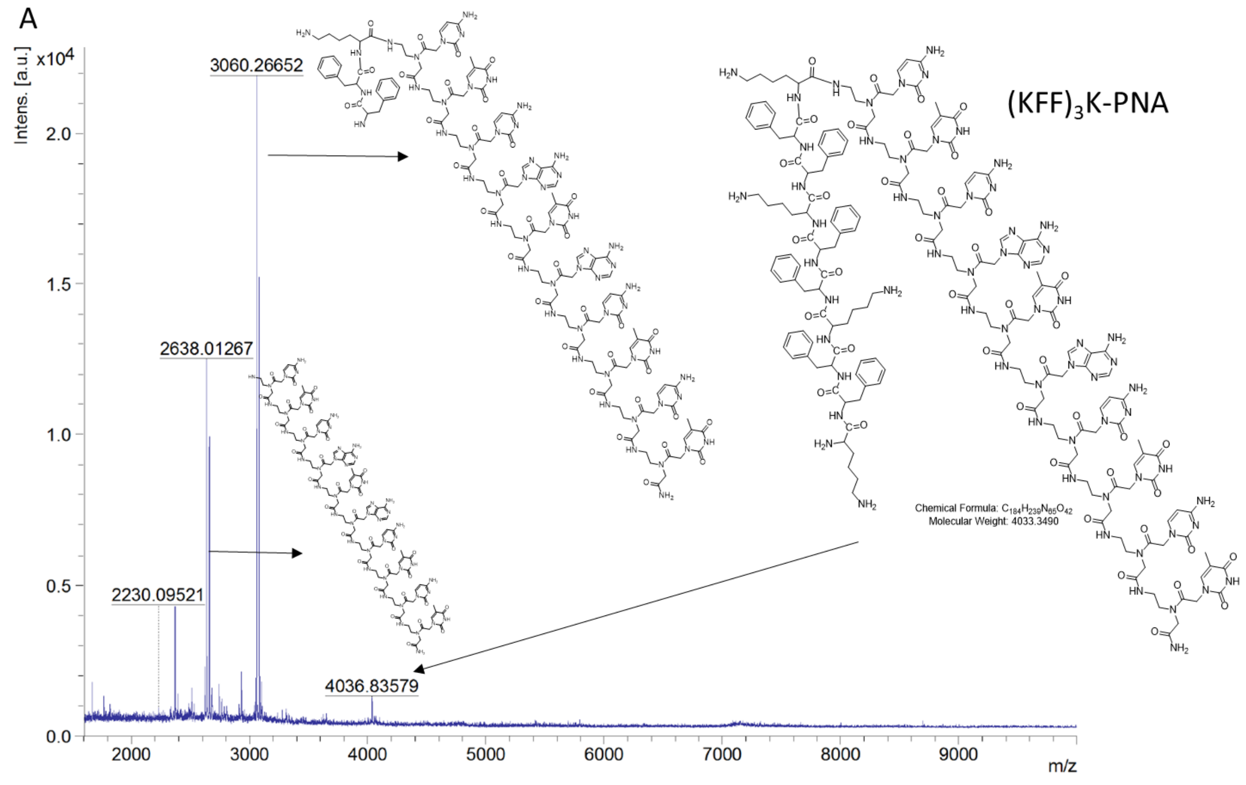


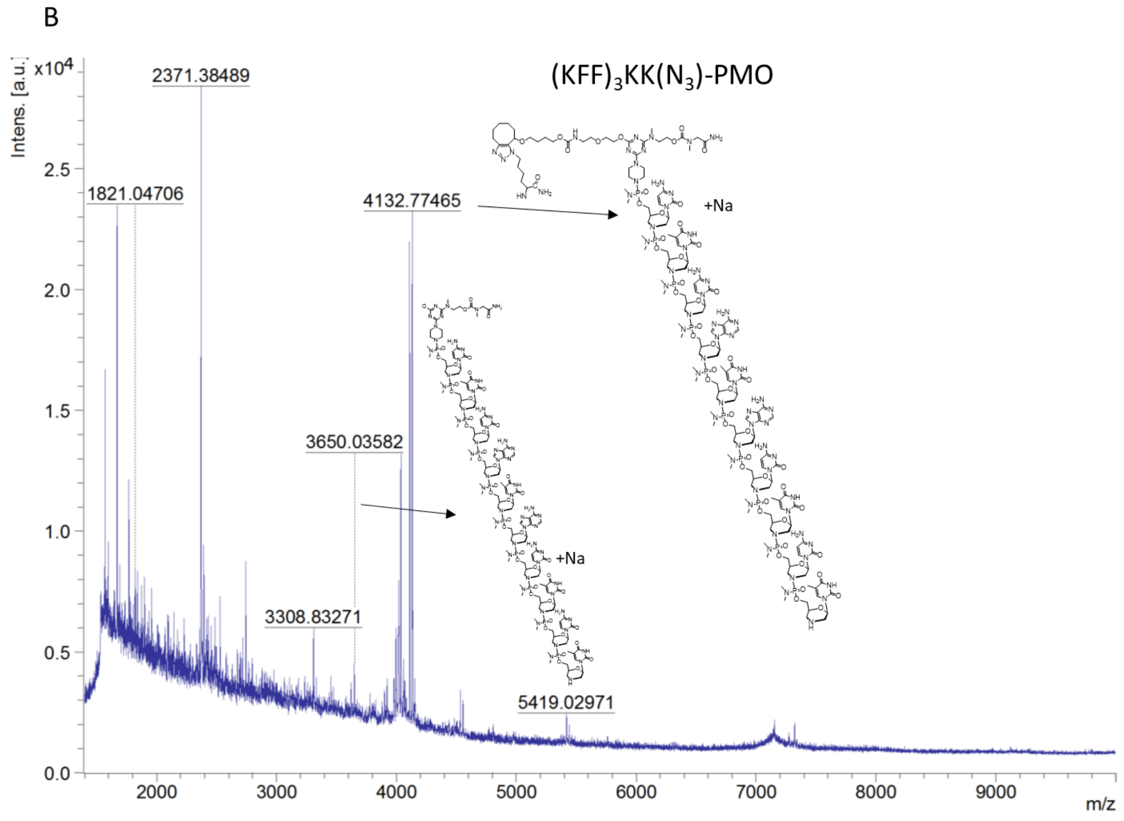


**Supplementary Figure S7.** MALDI-TOF based detection of fragments of KFF conjugates of PNA and PMO in periplasm of *Salmonella* 15 minutes post-treatment.

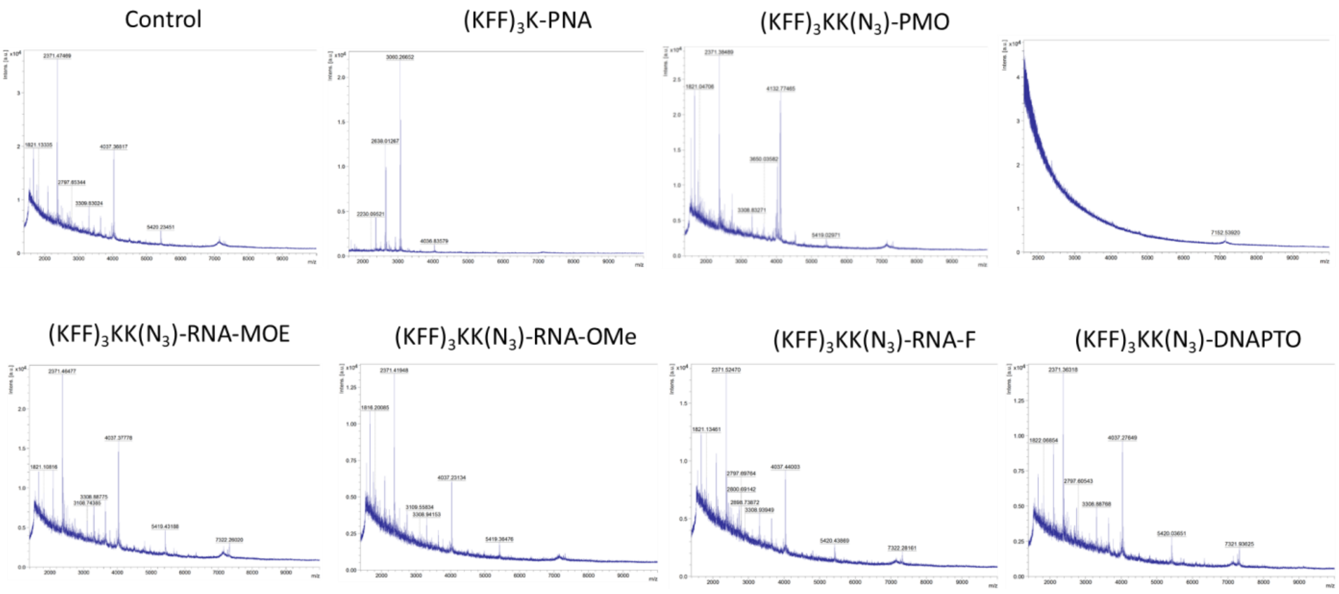


**Figure S8:** MALDI-TOF specta of periplasm of *Salmonella* post-treatment with different KFF-ASO conjugates.

**Table S1**. **Oligonucleotide sequences for generation of *gfp* fusion constructs.** Table includes internal oligonucleotide number (JVO-), target gene name, and the respective oligonucleotide sequence in 5' to 3' orientation. Fw: forward, rv: reverse. Underlined sequence: T7 promoter sequence, italic sequence: *gfp* overlap.

| **Oligo (JVO-)** | **Gene** | **Sequence (5´-3´)** |
|---|---|---|
| 19760 | acpP-fw | <u>TAATACGACTCACTATAG</u>AACCATCGCGAAAGCGAGTT |
| 19761 | acpP-rv | *TCCAGTGAAAAGTTCTTCTCCTTTGCTAGC*GCCCAGCTGTTCGCCGATAAT |
| 19762 | Gfp-fw | GCTAGCAAAGGAGAAGAACTTTTCAC |
| 19763 | Gfp-rv | TTATTTGTAGAGCTCATCCATGCC |
